# A platform for deep sequence-activity mapping and engineering antimicrobial peptides

**DOI:** 10.1101/2021.05.13.444096

**Authors:** Matthew P. DeJong, Seth C. Ritter, Katharina A. Fransen, Daniel T. Tresnak, Alexander W. Golinski, Benjamin J. Hackel

**Affiliations:** Department of Chemical Engineering and Materials Science, University of Minnesota – Twin Cities, Minneapolis, MN 55455, USA

**Keywords:** antimicrobial peptide, antibiotic, ribosome, oncocin, protein engineering, deep mutational scanning

## Abstract

Developing potent antimicrobials, and platforms for their study and engineering, is critical as antibiotic resistance grows. A high-throughput method to quantify antimicrobial peptide and protein (AMP) activity across a broad continuum can elucidate sequence-activity landscapes and identify potent mutants. We developed a platform to perform sequence-activity mapping of AMPs via depletion (SAMP-Dep): a bacterial host culture is transformed with an AMP mutant library, induced to express AMPs, grown, and deep sequenced to quantify mutant frequency. The slope of mutant growth rate versus induction level indicates potency. Using SAMP-Dep, we screened 170,000 mutants of oncocin, a proline-rich AMP, for intracellular activity against *Escherichia coli*. Clonal validation of 36 mutants supported SAMP-Dep sensitivity and accuracy. The efficiency and accuracy of SAMP-Dep enabled mapping the oncocin sequence-activity space with remarkable detail and scale and guided focused, successful synthetic peptide library design, yielding a mutant with two-fold enhancement in both intracellular and extracellular activity.

## Introduction

Antibiotic resistance is a critical global healthcare threat with 700,000 annual deaths and $55 billion in healthcare and productivity costs^1^ and an alarming upward trend.^2^ Along with improved antibiotic stewardship, innovative antimicrobial therapies and platforms for their efficient discovery are sorely needed. Antimicrobial peptides and proteins (AMPs) are a compelling alternative to traditional antibiotics.^3, 4^ AMPs kill bacteria via various processes and could be administered in combination to potentially reduce resistance.^5, 6^ Many AMPs are ribosomally synthesized, which permits efficient engineering of potency, specificity, and stability via recombinant DNA technology and innovative genotype-phenotype linkage strategies.^7, 8^ Therefore, rapid responsiveness is enabled to evolve AMPs as infectious microbes emerge.

Similar to ligand or enzyme engineering, high-throughput AMP engineering requires synthesis of a well-designed library and an effective sort, or screen, with accompanying high-throughput analysis to productively traverse immense sequence-function landscapes^9^. Precise, diverse mutagenic libraries are efficiently synthesized, and deep sequencing thoroughly samples large AMP sequence spaces, which motivates an efficient strategy to pair sequence to function for accelerated AMP development.

Innovative selection techniques have been developed for many functions, most powerfully for target binding^10, 11^ and organism survival^12, 13^. Yet AMPs pose a more complex, and thus challenging, set of mechanisms that ultimately result in cell death. Multiple methods have advanced AMP engineering and discovery. Multiwell plate assays afford limited throughput^14, 15^, which is only partially aided by recombinant AMP expression^16–19^. Innovative genotype-phenotype linkages, including surface-localized antimicrobial display (SLAY)^20^ and host-target co-encapsulation^21^, expanded throughput to 10^6^ variants, yet their current implementations provide a binary metric, active or inactive. Moreover, SLAY tethers the AMP to the cell surface, which limits mechanisms of action. Nevertheless, the concept that potent AMPs self-deplete via self-killing has been powerfully validated.

A binary metric is practical for discovery, but AMP engineering strategies would benefit from a continuous potency metric to differentiate a range of antimicrobial activities and map high-resolution sequence-activity landscapes. A recent study mutated α-synuclein, a growth-inhibiting protein, to deduce the active conformation in yeast.^22^ The sequence-activity landscape was mapped by tracking >10^3^ mutant frequencies after induction and scoring fitness as the slope of the log-transformed variant frequency over time. Baliga *et al*. coarsely screened 336 apidaecin AMP mutants with a similar approach.^23^ We present an enhanced approach to screen over 10^5^ AMP variants simultaneously, uncover sequence determinants of antimicrobial activity, and leverage the sequence-activity landscape to engineer AMPs with enhanced activity.

Oncocin was selected as the model antimicrobial because it is ribosomally synthesized, has intracellular targets, and is well characterized. First, oncocin, a 19- residue proline-rich peptide^24^, is encodable as a DNA template for expression within the host, enabling efficient screening of diverse libraries. Second, oncocin ablates bacterial growth by binding within the ribosome and inhibiting translation^25^, with the N-terminus deepest in the ribosomal exit tunnel, therefore, requiring post-translational ribosomal exit prior to inhibiting protein synthesis.^26, 27^ Third, there are known oncocin sequence-activity relationships, although moderate in breadth, that could inform library design and validate sequence-activity trends.^14, 26–29^

To efficiently map sequence-activity trends and engineer potent antimicrobials, we developed SAMP-Dep. We transformed a microbial host with a diverse AMP mutant library, induced intracellular AMP expression, and quantified depletion relative to induction level with deep sequencing to deduce activity. With SAMP-Dep, we engineered oncocin for enhanced activity against *E. coli*. The oncocin mechanism of action has been well-studied; however, to date, sequence-activity mapping has been limited to hundreds of mutants.^14^ To deeply map the functional landscape, a first-generation library sampled single and local double mutants, and an informed second-generation library evaluated broader multi-mutants. Through screening inducible variant depletion in a massively parallel fashion, SAMP-Dep provided a framework for deep, reproducible, and accurate sequence-activity mapping of over 170,000 oncocin mutants and guided focused peptide design with high activity rates and potency.

## Results

### SAMP-Dep was designed and tuned for high-throughput AMP engineering

The strategy for the high-throughput sequence-activity mapping SAMP-Dep platform is: first, the microbial host is transformed with an expression vector encoding an AMP variant library (**Figs. 1a,b**). The library of transformed hosts is induced at different strengths and grown. Cells expressing a less active AMP variant will grow more quickly than those harboring a more active variant (**Fig. 1c**). Growth rate is computed from deep-sequenced variant frequencies at each inducer level, and growth:induction slope is calculated as the potency score.

**Figure 1.**
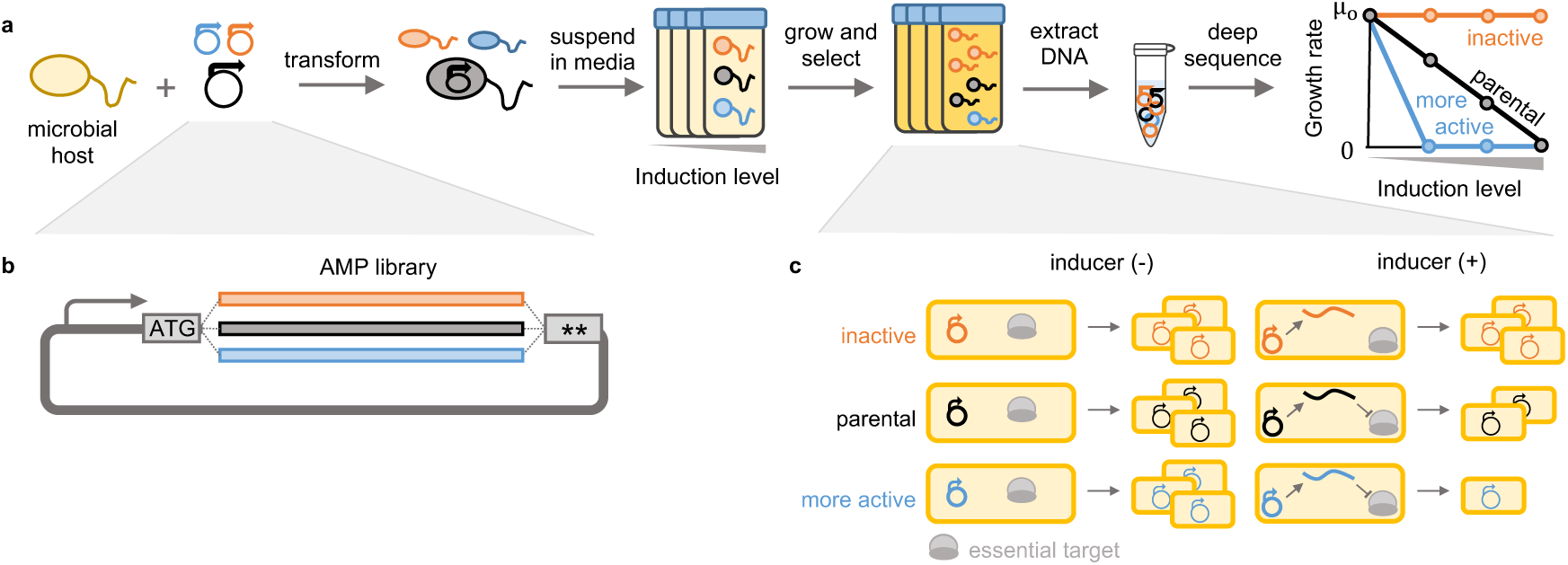
**SAMP-Dep quantified activity of AMPs with intracellular targets via deep sequencing. a**, Host microbes were transformed with a mutagenic AMP library and grown with varying levels of AMP induction. The resultant DNA was extracted from cells then sequenced, and clonal read frequencies were used to deduce antimicrobial activity. Potent AMPs were depleted relative to less functional AMPs at lower induction conditions. **b**, An expanded view of the mutagenic AMP library construct. **c**, All cells would grow without inducer. When AMP production was induced, inactive AMPs (orange) would grow uninhibited, the parental AMP (black) would slow growth, and more active variants (blue) would reduce growth further than parental.

We implemented SAMP-Dep with T7 Express LysY/I^q^ *E. coli* expressing oncocin mutants via an isopropyl β-D-1 thiogalactopyranoside (IPTG) inducible pET vector. The parental peptide, with a V1M (ATG) mutation to ensure translation, will be called oncocin in this paper. This system was selected because of tightly-controlled, inducible protein expression – to avoid depletion of potent mutants during library preparation – and the ability to move the system into alternative hosts/targets for broad use.^30–32^ *E. coli* harboring the oncocin expression vector effectively grew in the absence of IPTG inducer at a rate equivalent to a negative control (**Supplementary Fig. 1a**), confirming basal expression was sufficiently low to avoid growth inhibition. Oncocin expressing *E. coli* had an inducible reduction in growth from uninhibited growth at 0 mM IPTG to nearly complete growth inhibition at 0.50 mM IPTG, hypothetically enabling identification of enhanced antimicrobial variants at intermediate induction levels (**Figs. 1a,c** and **Supplementary Fig. 1a**).

Beyond validation of low basal expression and inducible growth inhibition, sampling time, inducer concentration, time of induction, and induction temperature for SAMP-Dep were determined in a series of pilot experiments (**Supplementary Figs. 1a-e**). With analysis via DNA sequencing, cell death was an insufficient outcome, for dead cell DNA could be sequenced; time was needed for viable cells to divide and achieve differential density. At least seven hours of growth allowed negative control accumulation and oncocin depletion (**Supplementary Fig. 1a**). The pilot experiment suggested 0.15 and 0.30 mM IPTG enabled partial depletion of oncocin while 0.50 mM IPTG enabled nearly full ablation of growth, enabling identification of a range of antimicrobial activities. Induction start time before or after transformation recovery showed negligible effect on culture density of cells expressing oncocin versus negative control while induction at 30 °C decreased the density difference relative to 37 °C (**Supplementary Figs. 1b-e**). Thus, we chose to induce AMP production at 0, 0.15, 0.30, and 0.50 mM IPTG at 37 °C after transformation recovery, and plasmid DNA would be extracted after at least seven hours of growth.

### First-generation library screened local single and double mutants

To systematically evaluate oncocin sequence-function relationships, a library of all double mutants at positions *n*/*n*+1 and *n*/*n*+2 was synthesized to provide full depth of single mutants and local double mutants while maintaining deep sequencing coverage (**Supplementary Fig 2a**). Due to oncocin binding linearly within the ribosome^26, 27^, pairwise interactions were expected to be primarily at neighboring amino acids, motivating the *n*/*n*+1 and *n*/*n*+2 design. NNN degenerate codons were included to assess synonymous mutation effect on potency. The mutations encompassed the entirety of oncocin, including the C-terminus, thought to be required for internalization^27^, to assess C-terminal determinants of intracellular activity.

Upon mutagenic gene assembly and bacterial transformation, sequencing revealed the desired diversity with the expected binomial distribution with one wrinkle: triple and quadruple codon mutations composed 5% of the library, likely from the incorporation of two mutagenic primers on a single PCR template (**Supplementary Fig. 2b**). Degenerate codons had a nearly equal split between A/T/G/C as desired (**Supplementary Fig. 2c**).

### SAMP-Dep quantified a range of antimicrobial activities

SAMP-Dep assayed activities of first-generation mutants: 9,237 peptide mutants within 27,480 DNA sequences including 8,740 of the 15,400 designed *n*/*n*+1 and *n*/*n*+2 amino acid variants. The remaining had three or four amino acid mutations. Across three replicates and four induction conditions, read counts were well correlated, indicating reproducibility (**Supplementary Fig. 2d**).

We computed the slope of growth rate versus inducer concentration as an activity metric because growth rate was independent of total mutant counts and approximately linear versus induction (**Fig. 2a**). Slope reproducibly quantified a continuum of antimicrobial activity (**Fig. 2b** and **Supplementary Fig. 2e**). Negative slopes, a decrease in growth rate when induced, suggested active variants. Slopes around zero, no change in growth rate, were likely non-functional. Consistent with expectation, oncocin had a moderately negative slope (-12 ± 2 M^-1^min^-1^), and truncated negative controls with nonsense mutations within the first ten positions had slopes of approximately 0 M^-1^min^-1^ (**Fig. 2b**).

**Figure 2.**
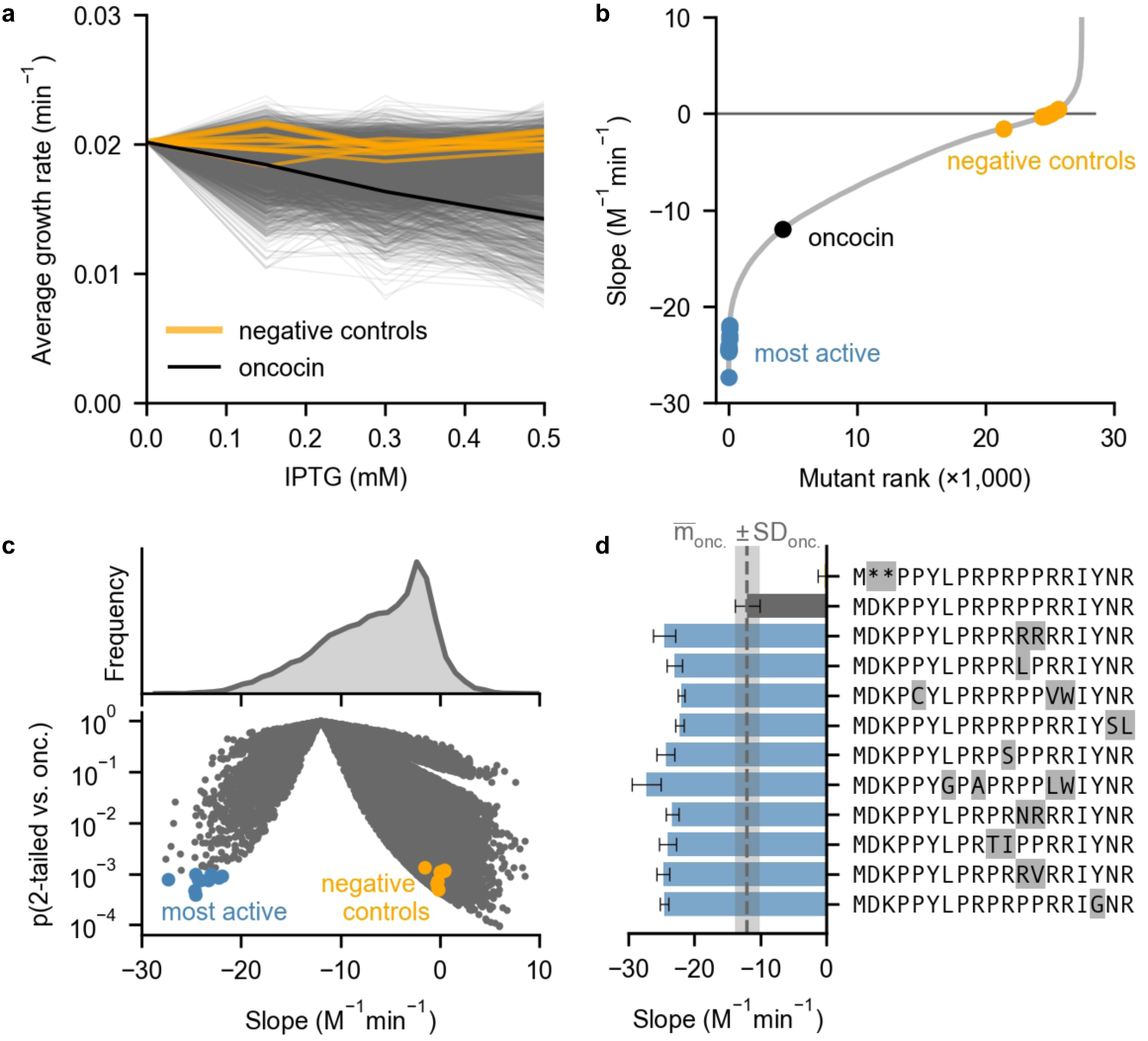
**SAMP-Dep identified oncocin variants with enhanced and diminished intracellular activity.** *Negative controls* indicate samples with nonsense mutations at sites 1-10 and error comparable to parental (slope standard deviation < 2 M^-1^min^-1^). **a**, Mutant growth rates varied with inducer concentration. **b**, First-generation oncocin mutants encoded a range of activities. **c**, Volcano plot of first-generation oncocin mutants revealed substantially more and less active oncocin variants. P value was calculated from a 2-tailed t-test relative to oncocin. **d**, Ten most potent mutants (blue) from Figure 2c (p < 5 × 10^-4^, 1-tailed t-test) relative to oncocin (grey) and a negative control. Error bars denote slope standard deviation from three SAMP-Dep replicates.

While most mutants lost function, ten mutants displayed enhanced intracellular activity relative to oncocin with p < 10^-3^ (1-tailed t-test), two of which contained more than two mutations (**Figs. 2c,d**). The most potent mutant, Y17G, was 2.0 ± 0.4 fold more inhibitory than oncocin (p = 2 × 10^-4^, 1-tailed t-test). Overall, SAMP-Dep paired over 27,000 DNA sequences to antimicrobial activity in a massively parallel fashion, deeply mapping the oncocin sequence-activity landscape.

### Second-generation library design leveraged first-generation relationships

The wealth of single and local double mutation sequence-activity relationships guided a second-generation library assessing broader multi-mutants. We hypothesized combinations of the most active mutations from the first generation could uncover potent multi-mutants. To extract potency information across all mutants, a linearly additive model trained on 80% of first-generation SAMP-Dep data accurately predicted the remaining 20%, indicating a sitewise-independent additive model could capture sequence-activity trends (**Supplementary Fig. 3a,b**). Mutations that broadly enhanced activity in the additive model (expected Δ slope < -1 M^-1^min^-1^) in addition to single mutants with comparable or enhanced activity relative to parental oncocin (slope < -9 M^-1^min^-1^) were incorporated in a second-generation library (**Fig. 3a** and **Supplementary Fig. 3b,c**). Internalization is critical for oncocin extracellular activity, so the second-generation library aimed to maintain internalization while enhancing intracellular activity. A previous study indicated a truncated oncocin mutant lacking the last seven residues (oncocin 1-12) lost extracellular activity while partially retaining *in vitro* translation inhibition.^27^ Acknowledging the C-terminus may affect internalization, little variation was introduced past P12 while positions 15-19 were preserved as parental (**Fig. 3a**). The library design did not incorporate premature stop codons, so negative controls encoding multiple early stop codons were included. The second-generation library design, which encoded 115,584 peptides within 161,664 codon-optimized DNA sequences including negative controls, captured nearly all desired mutant combinations with few exceptions due to library size and codon choice constraints.

**Figure 3.**
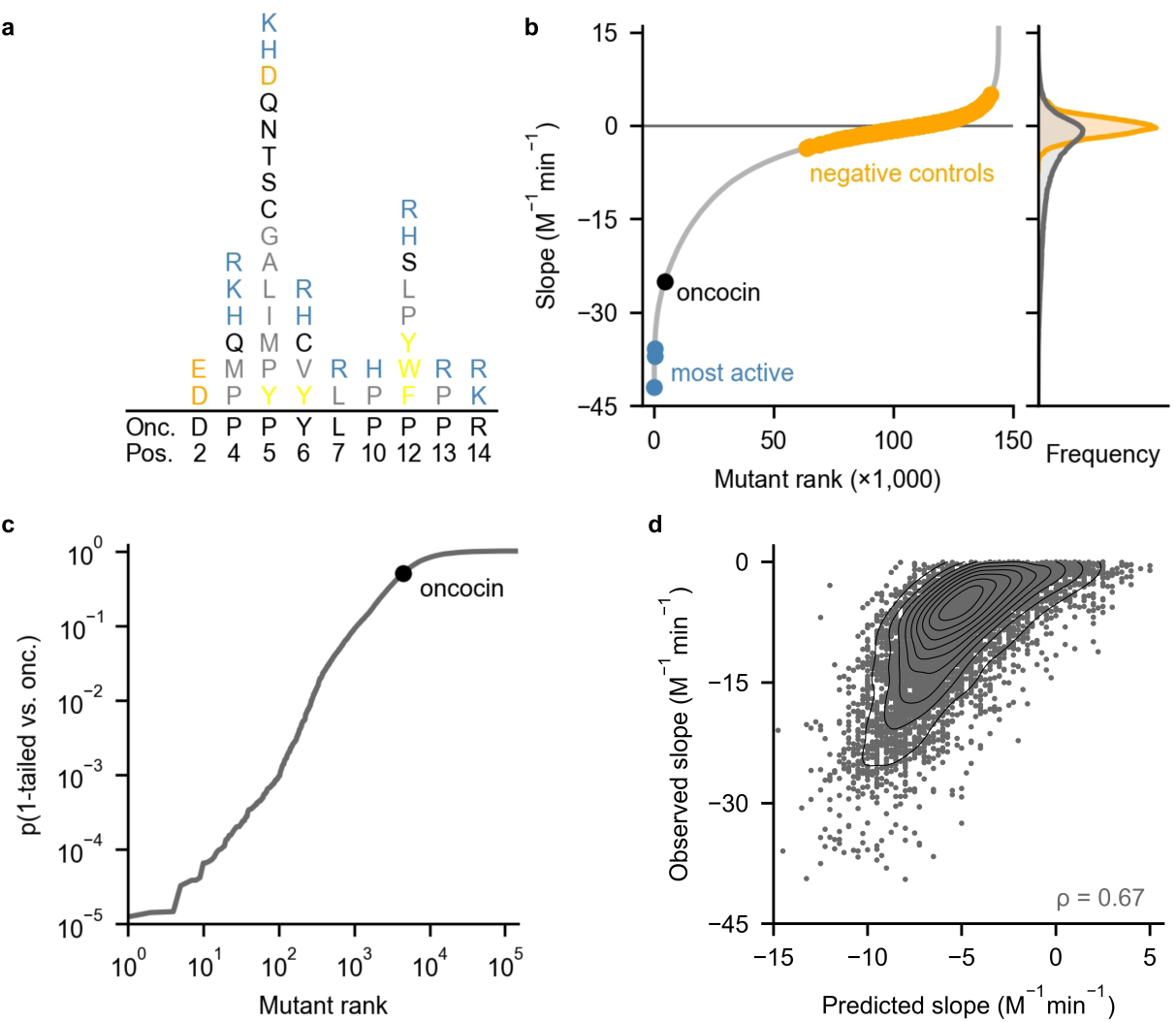
**SAMP-Dep found combinatorial oncocin mutations that enhanced activity. a**, The second-generation library was designed to contain mutants as random combinations of equal frequencies of the indicated amino acids (color denotes similar chemical properties) at the noted degenerate positions. **b**, *Left*: Growth:induction slope revealed a subset of highly active sequences among many inactive mutants. *Right*: slope histograms for all mutants (grey) and negative controls (orange). Negative controls were filtered for slope standard deviation < 2 M^-1^min^-1^. **c**, Many mutants displayed substantial improvements in activity relative to oncocin. P value was calculated from a 1-tailed t-test relative to oncocin. **d**, An additive model trained on 80% of second-generation sequence-slope pairs predicted slope for the remaining 20% of second-generation sequences. All mutants were filtered for slope standard deviation < 2 M^-1^min^-1^.

SAMP-Dep assayed second-generation multi-mutant activities via induction at 0, 0.10, 0.20, and 0.50 mM IPTG (**Supplementary Fig. 3d**). The 143,994 unique DNA sequences screened 89% and 94% of desired DNA and amino acid diversity, respectively. Growth:induction slope reproducibly mapped the second-generation sequence-activity landscape (**Fig. 3b** and **Supplementary Figs. 4a,b**). 63 mutants displayed substantially enhanced activity (p < 5 × 10^-4^, 1-tailed t-test relative to oncocin), while only ten mutants were as enhanced in the first generation (**Fig. 2d** and **Fig. 3c**). Second-generation mutants with enhanced activity were more sequence-similar to oncocin (mean number of mutations = 4.8) relative to the entire library (mean = 6.1) (p < 0.05, χ^!^), suggesting active mutants occupied a subset of the sampled sequence space (**Supplementary Fig. 4c**).

The first and second-generation libraries had an overlap in sequence space, enabling comparison. Their growth:induction slopes were correlated, but second-generation slopes were typically (15 of 16) more negative than first-generation, likely due to different parental DNA sequence (**Supplementary Fig. 4d**). Aligned with the observed correlation, the first-generation additive model was predictive of second-generation activity, suggesting the model captured general sequence-activity relationships (**Supplementary Fig. 4e**). The first-generation model predicted second-generation mutants that followed the first-generation *n*/*n*+1 and *n*/*n*+2 design better than broader second-generation multi-mutants, suggesting the discrepancy was related to higher-order effects not captured by the linearly additive model used in designing the second-generation library (**Supplementary Fig. 4f**). An additive model trained on 80% of second-generation SAMP-Dep data predicted activity of the remaining 20%, further demonstrating the utility of additive models for predicting oncocin activity (**Fig. 3d**).

### SAMP-Dep identified AMPs with a range of antimicrobial activity

Mutants were clonally validated to assess if SAMP-Dep accurately captured intracellular activity at library scale. Accuracy was systematically evaluated via four questions: 1) Was SAMP-Dep growth:induction slope at library scale sensitive to intracellular activity? 2) Were mutants with substantially decreased slope more intracellularly active? 3) To what extent did codon choice affect activity? 4) Did mutants with positive slope increase growth rate with induction? (**Supplementary Figs. 5a,b**).

SAMP-Dep slope measured at library scale was sensitive to intracellular activity. Individual mutant growth rates were directly measured in T7 LysY/I^q^ *E. coli*, and a clonal growth:induction slope was quantified for 36 mutants with a range of SAMP-Dep slope and low slope error (slope standard deviation ≤ 6 M^-1^min^-1^). SAMP-Dep slope broadly correlated with clonal slope for first and second-generation mutants (ρ = 0.81; **Fig. 4a**). The strong correlation of SAMP-Dep slope with clonal slope suggested SAMP-Dep uncovered genuine sequence-activity relationships in high-throughput.

**Figure 4.**
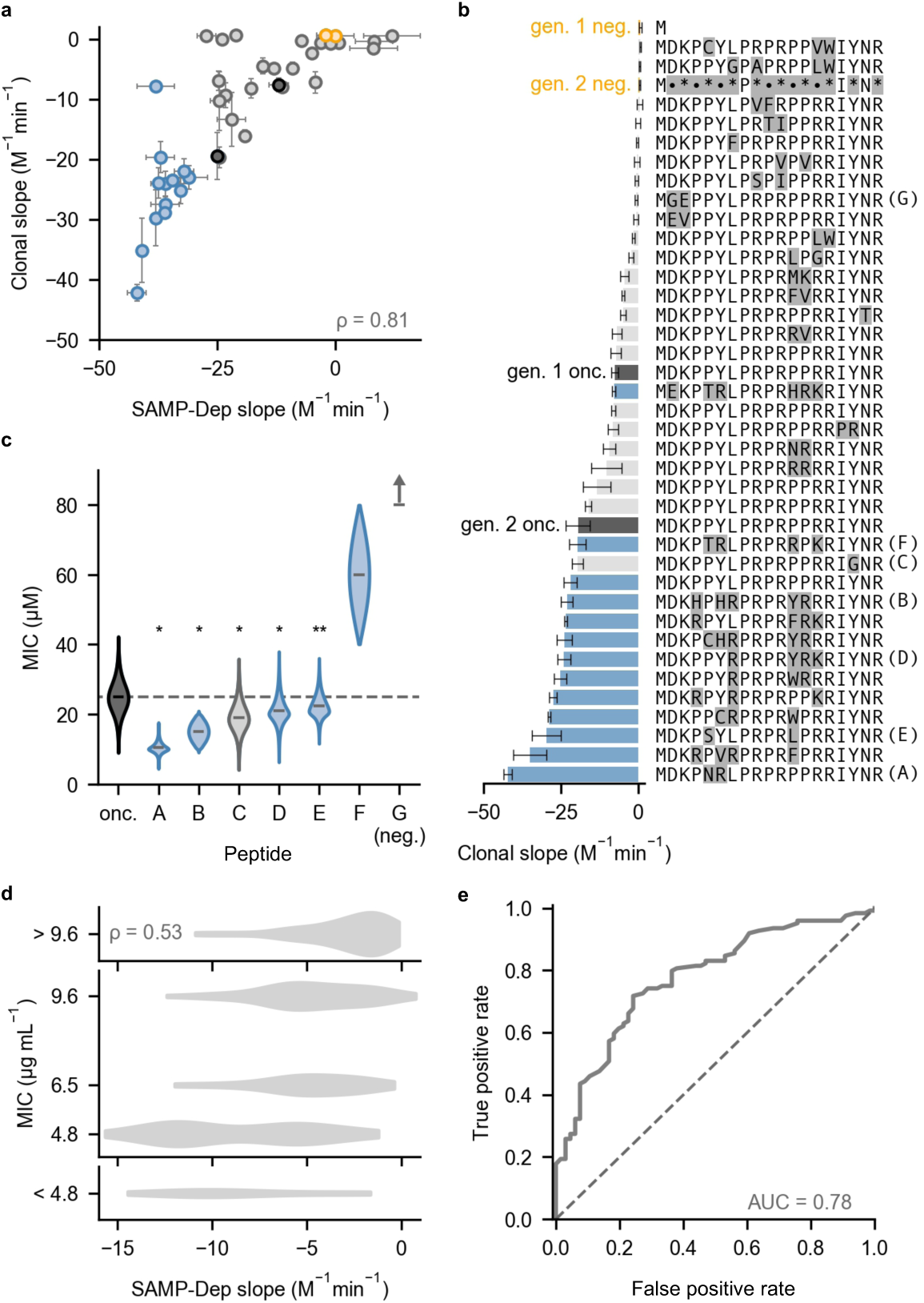
**SAMP-Dep enhanced intracellular and extracellular activity. a**, Clonal growth:induction slope was well correlated with SAMP-Dep slope for 36 mutants from Gen. 1 (grey) and Gen. 2 (blue). Negative controls (orange) and oncocin (black) were assessed from both generations. Points represent mean, and the error bars represent slope standard deviation from three replicates for both SAMP-Dep and clonal experiments. DNA sequences with corresponding SAMP-Dep and clonal slopes and errors were tabulated in **Supplementary Table 1**. A cluster of three first-generation mutants was expected to enhance activity but was inactive. One second-generation mutant expected to enhance activity did not, but the mutant retained clonal activity comparable to first-generation oncocin. Some false positives were expected given the high-throughput nature of SAMP-Dep. Deviation from the correlation was weakly correlated with both SAMP-Dep slope error and uninduced read count but did not completely explain the false positive (**Supplementary Figs. 5c,d**). **b**, Clonal mutants had a range of intracellular activities. Bars represent mean, and the error bars represent slope standard deviation from three SAMP-Dep replicates. • indicates a degenerate position for second-generation negative control. Colors represent the same categories as Figure 4a. **c**, Synthetic peptides with enhanced SAMP-Dep activity displayed antimicrobial activity against T7 Express LysY/I^q^ host. * indicates p < 10^-5^, and ** indicates p < 4 × 10^-3^ from bootstrapping relative to oncocin. Negative control peptide G did not inhibit growth at 80 µM. Colors represent the same categories as Figure 4a. **d, e**, SAMP-Dep slope of 190 variants compared to the literature precedent^14^ for extracellular activity against *E. coli* JW0013, a DnaK knockout strain. **e**, Receiver operating curve using SAMP-Dep slope as the classifier variable (solid) relative to a random classifier (dashed) to predict peptide extracellular activity. Peptides with MIC ≤ 9.6 µg mL^-1^ were deemed active.

Clonal mutants with enhanced SAMP-Dep slope were typically more active. Of the 36 clonal mutants, 7 of 12 first-generation mutants and 12 of 13 second-generation mutants with enhanced SAMP-Dep slope relative to parental oncocin were also more active in the clonal assay (**Fig. 4b**). Oncocin Y17G, the most potent mutant from the first generation, was the most potent clonal mutant from the first generation with a 2.6 ± 0.6 fold enhanced clonal activity relative to first-generation oncocin. Oncocin P5N Y6R, the most potent mutant from the second generation, was the most potent clonal mutant quantified with a 2.2 ± 0.4 fold enhanced clonal activity relative to second-generation oncocin (**Fig. 4b** and **Supplementary Table 1**). Although identified in the second generation, the P5N Y6R mutation fit the first-generation library design: oncocin P5N Y6R was one of the most potent first-generation variants with a growth:induction slope of -18.9 ± 0.9 (p = 2 × 10^-3^, 1-tailed t-test relative to oncocin), consistent with expectation. SAMP- Dep successfully identified oncocin mutants with enhanced intracellular activity.

Codon choice influenced activity of synonymous mutants. Five of the 36 clonal mutants were synonymous with parental oncocin, quantifying transcript effect on activity. Between generations, the second-generation parental oncocin sequence displayed a 2.4 ± 0.6 fold enhancement in clonal activity relative to first-generation oncocin, suggesting the first-generation transcript was less producible (**Fig. 4b**). Synonymous oncocin mutants within the clonal validation library enabled further comparison (**Supplementary Fig. 5a**). The putative decreased activity first-generation mutant encoding oncocin had 0.9 ± 0.3 fold reduced activity, while the putative enhanced variant had a 1.8 ± 0.6 fold improved clonal activity (**Fig. 4b**). Most notably, the second most potent first-generation clonal variant was a synonymous oncocin mutant with 2.1 ± 0.3 fold enhanced activity, nearing the activity of second-generation parental oncocin. For the second-generation clonal library, one synonymous oncocin mutant was expected to enhance activity (**Supplementary Fig. 5b**). The synonymous mutant had comparable activity to second-generation parental oncocin (1.1 ± 0.2 fold, **Fig. 4b**). Variance among mutants encoding parental oncocin indicate the role of codon choice in perceived activity, especially within the first generation. Mutants with nominally positive SAMP-Dep slopes were inactive in clonal testing, which was reasonable given their high SAMP-Dep slope error (**Fig. 4a** and **Supplementary Fig. 5a**). Taken together, clonal validation suggested SAMP-Dep accurately quantified intracellular activity in a massively parallel fashion.

Oncocin extracellular activity is conditional on intracellular activity; therefore, SAMP-Dep slope was expected to be partially predictive of extracellular activity. The most potent first-generation mutant (2.0 ± 0.4 fold enhancement relative to parental) along with five potent second-generation mutants (ranging from 1.2 to 1.6 fold enhancement relative to parental) were selected for exogenous validation (**Supplementary Table 1)**. A full-length mutant with a SAMP-Dep slope of 0.8 ± 0.5 M^-1^min^-1^, D2G K3E, was included as a negative control. Synthetic peptides were administered to T7 LysY/I^q^ *E. coli* to measure a minimal inhibitory concentration (MIC). Parental oncocin exhibited a MIC of 25 ± 5 µM, which agreed with previously quantified^27^ 25 µM MIC for Onc112 against *E. coli* BL21, the unmodified strain of T7 LysY/I^q^ *E. coli*. The negative control peptide was inactive at 80 µM, the highest concentration assayed (**Fig. 4c**). All mutants with enhanced clonal activity had observable exogenous activity, and five of six validated mutants had moderately enhanced activity relative to oncocin (p < 4 × 10^-3^, bootstrapping relative to oncocin; **Fig. 4b,c**). Peptides A and B had substantial improvements in exogenous activity, exhibiting 2.4 ± 0.6 and 1.7 ± 0.4 fold enhancement (p < 10^-5^, bootstrapping relative to oncocin), respectively (**Fig. 4c**). Notably, the DNA sequence encoding peptide A, corresponding to oncocin P5N Y6R, was the most active variant observed in both SAMP-Dep and clonal experiments (**Fig. 4a,b**). Therefore, SAMP-Dep successfully engineered oncocin multi-mutants with enhanced activity.

A previously published synthetic peptide library of site-saturated monosubstituted oncocin mutants from positions 1-11 against *E. coli* JW0013, a DnaK knockout strain, enabled independent comparison of 190 first-generation mutant SAMP-Dep slopes to literature^14^ MIC (**Fig. 4d**). All seven synthetic peptides with SAMP-Dep slope less than oncocin had observable activity, six of which had equal or enhanced MIC. A set of seven random peptides would have a 5% chance to be all active (MIC ≤ 9.6 µg mL^-1^) and only a 0.06% chance that at least six also had equal or greater activity. Further, SAMP-Dep slope was predictive as a binary classifier for extracellular activity (receiver-operator curve area of 0.78, **Fig. 4e**). Comparison with literature precedent supported the notion that screening of intracellular activity enhances hit rates of guided synthetic peptide libraries.

### Deep sequence-activity landscape mapping revealed trends in activity

The oncocin sequence-activity landscape characterized by SAMP-Dep agreed with known relationships (**Fig. 5a**). The PRP signature motif of proline-rich homologs was conserved at positions 8-10 within functional mutants, reflecting the importance of the PRP motif for intracellular activity.^27, 33^ Positions D2, K3, and R11, residues with interactions critical for ribosomal binding^25–27^, were intolerant to mutation. In the co-crystal structure of wild-type oncocin binding to the ribosome, P5 displaces ribosomal nucleotide U2585^26^, suggesting the observed mutational tolerance may relieve steric hinderance. Additionally, aromatic amino acid substitutions to P12 enhance activity against *Staphylococcus aureus*^34^, consistent with observed mutational tolerance of P12 to aromatic groups. Overall, consistency with known relationships suggested SAMP-Dep quantified genuine sequence-activity relationships.

**Figure 5.**
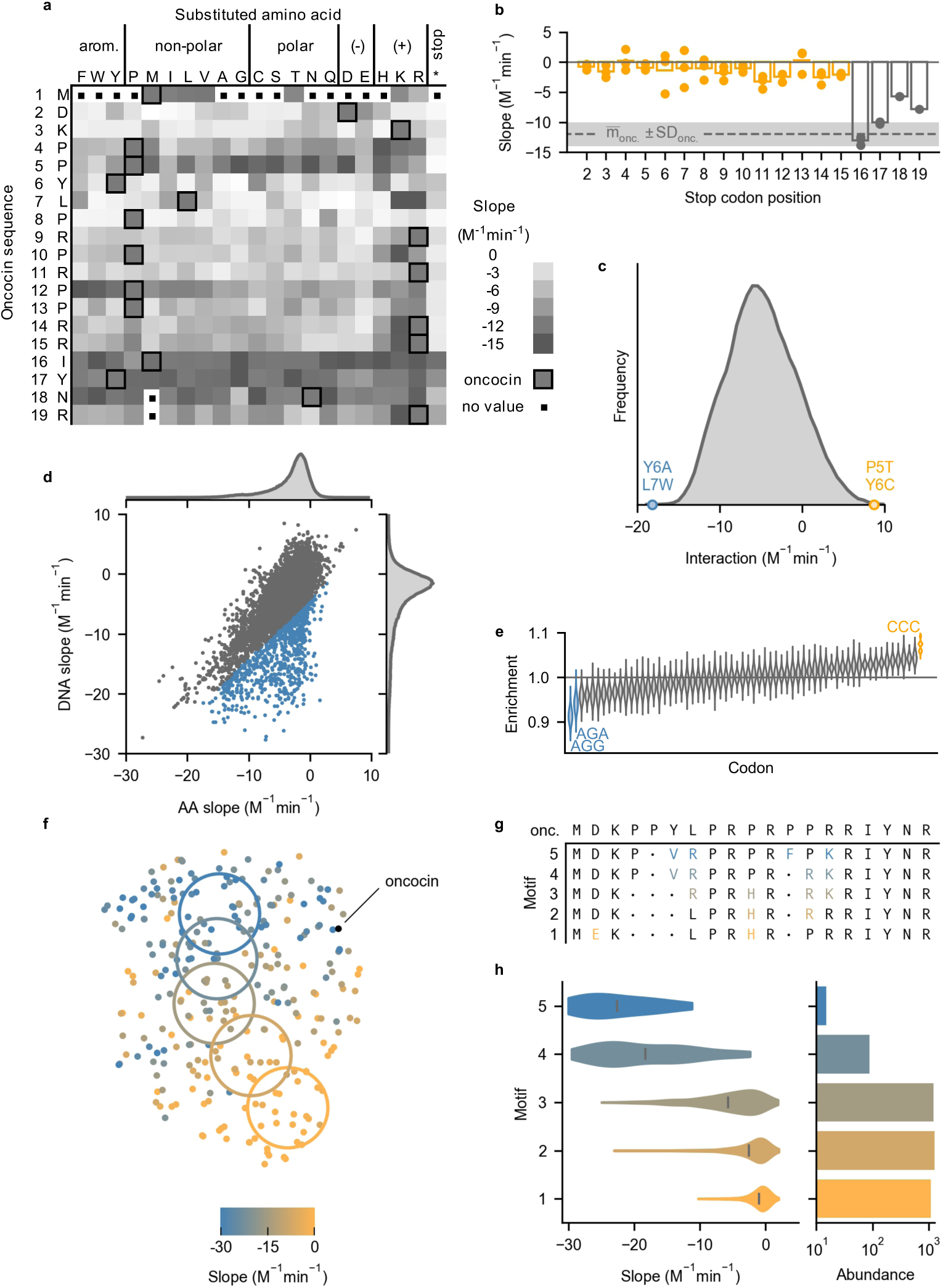
**SAMP-Dep revealed oncocin sequence-activity relationships.** All slopes represent growth:induction SAMP-Dep slope. **a**, Single mutant slopes from the first generation revealed trends in activity. Darker squares represent higher activity. Oncocin amino acids are outlined at each site. A distribution of slope error is included in **Supplementary Figure 6b**. **b**. A restricted heatmap of mutants with slope standard deviation less than 4 M^-1^min^-1^ from three replicates is included in **Supplementary Figure 6c**. **b**, Premature stop mutants regained activity past position 16. Bar and points represent the average and individual slopes of mutants with the first stop at the indicated position. The dashed line and shaded region indicate parental slope and standard deviation. **c**, Double mutant cycling of growth:induction slope revealed double mutants Y6A L7W (blue) and P5T Y6C (orange) with substantial epistasis. Interaction = Δslope_mut 1&2_ - Δslope_mut 1_ - Δslope_mut 2_. Mutants were filtered for slope standard deviation < 2 M^-1^min^-1^. Interaction is presented with kernel smoothing. **d**, The slope of each DNA sequence is plotted versus the average slope of all synonymous peptide sequences. **e**, Synonymous mutations to a CCC codon decreased intracellular activity while mutation to AGG and AGA improved activity. Enrichment was the frequency of codons in the grey region relative to overall frequency in **d**. Distributions were calculated via resampling with replacement. **f**, T-distributed stochastic neighbor embedding of 291 one-hot encoded AMP amino acid sequences revealed a gradient of intracellular activity across sequence space. Mutants were filtered for SAMP-Dep slope standard deviation less than 2 M^-1^min^-1^ and randomly sampled to form a uniform SAMP-Dep slope distribution. **g**, Consensus mutations (individual mutations present in greater than 50% of mutants) displayed higher-order sequence-activity relationships. Motif mutation color corresponded to the colored region in Figure 5f. • indicates no consensus mutation. **h**, *Left*: Second-generation mutant slope distribution with motifs identified in Figure 5g. *Right*: Motif prevalence in the second-generation library.

The oncocin sequence-activity landscape also agreed with active homologs with conserved ribosomal binding conformations^28^ (**Supplementary Fig. 6a**). Oncocin P5D L7R, a metalnikowin analog, had a slope of -20 ± 1 M^-1^min^-1^, relative to an oncocin slope of -12 ± 2 M^-1^min^-1^, and L7R commonly maintained activity in oncocin mutants (**Fig. 4b**). Oncocin P4G P5S, a pyrrhocoricin analog (past site 11, the binding conformation diverges^28^), had a slope of -9 ± 3 M^-1^min^-1^. Bac7 is identical to oncocin at 8 of 9 sites in the proline-rich region with the mismatch (oncocin Y6 / bac7 R9) occupying the same ribosome binding site.^28^ Four synonymous DNA mutants encoding Y6R had slopes of - 16 to -20 M^-1^min^-1^, while a fifth mutant had a slope of -3 ± 8 M^-1^min^-1^. The most potent mutant discovered, oncocin P5N Y6R, also contained the bac7 homolog mutation (**Figs. 4b,c**).

Building upon known sequence-function relationships, SAMP-Dep provided insight to potential sequence determinants of intracellular oncocin activity. Previous structural studies suggested the C-terminal segment past position P12 was not involved in ribosomal binding.^26, 27^ Recently, intracellular expression of oncocin revealed C-terminus truncated peptides retained activity until truncation at position 15.^29^ In agreement, our results suggest residues 1-15 are required for intracellular activity, not positions 1-12 as hypothesized by structural studies^27, 28^, for any truncation before position 16 broke intracellular activity, as indicated by slope around 0 M^-1^min^-1^ (**Fig. 5b**). In fact, positions R14 and R15 were intolerant to mutation, further suggesting their importance for intracellular activity (**Fig. 5a**). Tolerance to mutation, or truncation, at positions 16, 17, 18, and 19 indicated they are not likely involved with intracellular activity.

Double mutation cycling, comparison of the sum of slope changes for two separate mutations relative to the slope change for the double mutant, systematically revealed epistasis. A notable epistatic interaction was the Y6A L7W mutational pair, with an interaction of -18 ± 3 M^-1^min^-1^, indicating the double mutant was substantially more potent than expected under additivity (**Fig. 5c**). Y6 is important for ribosomal binding, for Y6A weakens ribosomal binding affinity and minimal inhibitory concentration by over 7 and 16 fold, respectively.^25, 35^ Additionally, Y6 and L7 have a favorable stacking interaction with the 70S ribosome^26, 27^, so Y6 and L7 were perhaps primed for epistasis. Y6A and L7W single mutants were essentially inactive with slopes of -0.9 ± 0.7 and -0.7 ± 0.4 M^-1^min^- 1^, respectively; yet the Y6A L7W double mutant regained activity with a slope of -8 ± 2 M^- 1^min^-1^. Further experimentation would be necessary to understand whether site 7 tryptophan, an aromatic residue capable of π stacking and hydrogen bonding like tyrosine, functionally replaced Y6, which forms a π stacking interaction with ribosomal nucleotide C2452^26, 27^ and hydrogen bonds with an unidentified ion coordinated by ribosomal nucleotides^27^. Alternatively, P5T Y6C exhibited clear epistasis with an interaction of 9 ± 3 M^-1^min^-1^, suggesting P5T Y6C was less active than expected. The *n*/*n*+1 *n*/*n*+2 library design enabled discovery of the Y6A L7W and P5T Y6C epistatic pairs via double mutation cycling.

NNN degenerate codons in first-generation library construction quantified codon choice effect in synonymous mutations. SAMP-Dep slope for particular DNA sequences relative to the average of synonymous amino acid sequences exhibited substantial correlation (**Fig. 5d**), consistent with a lack of codon preference. Yet, some variants differed from this correlation; analysis of codon usage in such deviants revealed that the CCC proline codon (p < 2 × 10^-4^, bootstrapping), rare in *E. coli*, decreased activity while the rare AGG and AGA arginine codons enhanced activity (p < 2 × 10^-4^ for AGG and p = 1.4 × 10^-3^ for AGA, bootstrapping) (**Fig. 5e**).^36^ The most potent synonymous oncocin mutant had two codon mutations: CGT → CGG and CGC → AGG, further implicating a role of AGG in first-generation transcript activity (**Fig. 4b**). The parental oncocin sequence contained two CCC codons but lacked AGG and AGA codons, suggesting the CCC tRNA was depleted while the AGG and AGA tRNAs were perhaps underutilized. Synonymous mutations highlighted how codon choice could impact potency, and, thus, DNA sequence effects should be considered when interpreting SAMP-Dep sequence-activity relationships. The dominant correlation of DNA and amino acid sequences highlights that this effect was often small.

In the second generation, SAMP-Dep characterized useful features of higher-order sequence space (**Figs. 3a,b**). To understand higher-order sequence determinants of antimicrobial activity, mutant amino acid sequences were visualized in two dimensions via one-hot encoding followed by t-distributed stochastic neighbor embedding (t-SNE)^37^, revealing a gradient of activity (**Fig. 5f**). Systematically sampling mutants across the two-dimensional space along the gradient of activity identified higher-order sequence determinants of activity (**Figs. 5f,g**). D2E was a consensus mutation among the non-functional mutant cluster, suggesting its role in ablating activity. D2E has been shown to reduce intracellular and extracellular activity^27^, and the least active clonal mutants validated from each generation had a D2E mutation (**Fig. 4b**). Traversing clusters towards higher activity, P10H was depleted, P13R was enriched then depleted, and L7R and R14K were introduced (**Fig. 5g**). The most potent mutant cluster displayed consensus mutations Y6V L7R P12F R14K. Oncocin P4R Y6V L7R P12F was the second most potent mutant clonally validated and contained part of the consensus motif (**Fig. 4b**). More broadly, second-generation mutants encoding the identified motifs displayed a similar gradient of SAMP-Dep activity (**Fig. 5h**). Other trajectories revealed small motif variations (**Supplementary Figs. 6d-f**). The t-SNE representation of sequence space supported the notion that P5 and P12 were tolerant to second-generation mutations as suggested by the first-generation (**Fig. 3a**, **Figs. 5a,g**, and **Supplementary Fig. 6f**). The higher-order sequence-activity relationships discovered by SAMP-Dep highlight the utility of deeply mapping sequence-activity space to uncover uncharted functional sequence space practically unattainable through traditional screening methods.

### Determinants of SAMP-Dep performance

To understand how experimental design influenced SAMP-Dep accuracy and precision, we systematically removed replicates, induction conditions, and read counts and assessed reduction in performance relative to the complete dataset (all three replicates, all induction conditions, and full sequencing depth). We calculated sensitivity and precision for identifying mutants significantly more active than oncocin (p < 0.05, 1-tailed t-test). All replicates and most induction conditions were beneficial to SAMP-Dep robustness (**Figs. 6a,b**). Across both first and second-generations, triplicates provided substantial benefit (**Fig. 6a**). The full induction condition, 0.50 mM IPTG, was critical for SAMP-Dep robustness, while the higher intermediate induction conditions added value (**Fig. 6b**). Lower intermediate induction conditions did not substantially aid performance. Our data demonstrates the importance of performing multiple replicates at several induction conditions, particularly higher inducer concentrations.

**Figure 6.**
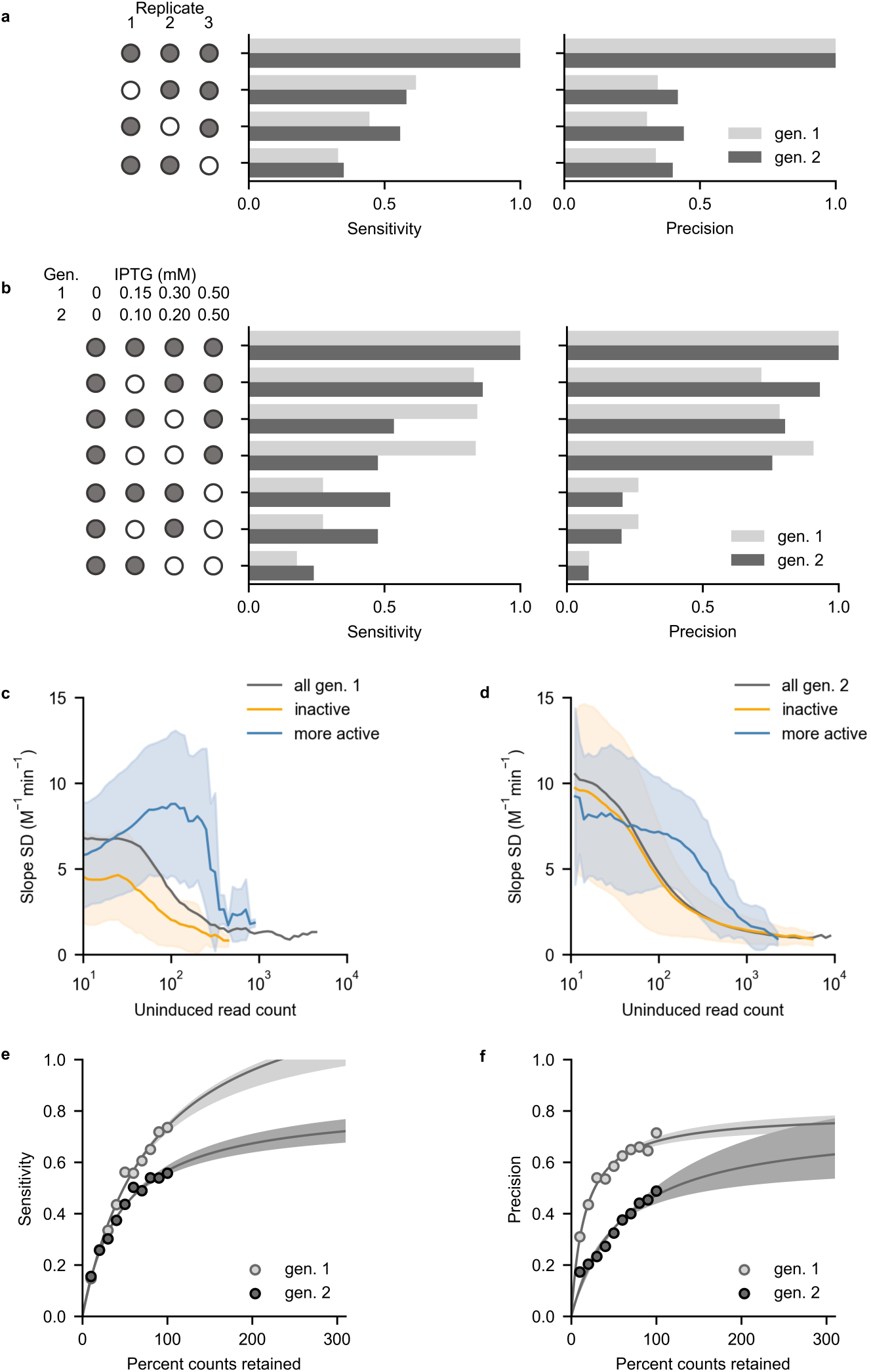
**Determinants of SAMP-Dep accuracy and precision.** Mutants that had significantly more negative slope than oncocin (p < 0.05 via a 1-tailed t-test) were defined as positive. **a**, Fraction of total true positive variants identified (sensitivity) and fraction of true positives among classified positives (precision) benefited from all three replicates. Filled circles indicate included replicates. **b**, Full inducer concentration, 0.50 mM IPTG, substantially benefited sensitivity and precision, and intermediate inducer concentrations also added value. Filled circles indicate included induction conditions. **c and d**, Slope standard deviation converged on 2 M^-1^min^-1^ as average uninduced read count approached 10^3^ counts for variants with nominally enhanced activity in the first (**c**) and second generations (**d**). Less potent slope error converged at lower read counts. The line and shaded region indicate the average and standard deviation of slope within the uninduced count bin. **e**, Sensitivity increased with simulated percent counts retained for the first generation (light grey) and second generation (dark grey). Shaded regions indicate a 95% confidence interval (F-test) about the nonlinear fit. **f**, Precision increased with simulated percent of original counts retained for the first generation (light grey) and second generation (dark grey). Shaded regions indicate a 95% confidence interval (F-test) about the nonlinear fit.

Sequencing depth aided SAMP-Dep accuracy and precision. Abundant mutants had more reproducible SAMP-Dep slope (**Supplementary Figs. 7a,b,c**). More potent mutants required higher uninduced prevalence for consistently reproducible SAMP-Dep slope relative to less active variants, suggesting increasing depth would aid quantification of active variants (**Figs. 6c,d**). Stochastically reducing sequencing depth decreased both the sensitivity and precision of identifying substantially enhanced mutants (**Figs. 6e,f**). Extrapolation from the non-linear fit suggested both sensitivity and precision could be improved with deeper sequencing, although the first-generation precision appeared to plateau. The analysis suggested increased replicates, higher induction, and increased sequencing depth aid activity quantification.

## Discussion

Broad and deep evaluation of immense protein sequence spaces benefits therapeutic discovery and fundamental elucidation of AMP sequence-activity relationships. Previous sequence-activity mapping has been hindered by complex antimicrobial phenotypes, traditionally screening hundreds of AMP variants at most. The efficient and accurate bandwidth of SAMP-Dep enabled, to our knowledge, the most comprehensive sequence-activity mapping of an AMP to date. 170,000 oncocin variants were evaluated in a massively parallel fashion, which enabled efficient engineering of oncocin for enhanced antimicrobial activity when administered both endogenously and exogenously. Moreover, cellular transformation and deep sequencing techniques readily enable SAMP-Dep to assess millions of variants. The current work complements the SLAY^20^ platform to further develop the concept of AMP discovery or engineering via deep sequencing to reveal depletion of highly potent AMP genes acting upon their host cell. SAMP-Dep’s ability to map sequence-activity relationships for intracellular targets complements the solely extracellular targeting of SLAY. SAMP-Dep’s use of soluble, untagged AMPs is also distinct from SLAY’s tethered system.

SAMP-Dep probes intracellular activity, which can be modulated by target affinity, target specificity, proteolytic stability, producibility, or a different trait. Simultaneous evaluation of these mutually important elements aids efficiency yet also requires deconvolution if a particular enhanced mechanism is desired. Other techniques could assess biophysical metrics including stability via deep sequencing of flow cytometrically sorted yeast displayed protein exposed to protease^38, 39^ and expression via a library-scale split green fluorescent protein assay^39^ or dot blots for mid-throughput.

The SAMP-Dep platform should be readily extended to alternative cellular hosts/targets. The key technological hurdles will be to achieve high transformation for library breadth and tight transcriptional control to assess uninduced variant levels. Non-leaky, inducible control of AMP expression has been achieved in pBAD systems in *E. coli*, not requiring a T7 RNA polymerase.^23, 29^ Similarly, extension to AMPs beyond oncocin is conceptually straightforward with the current limitation of requiring an intracellular target, which encompasses a broad swath of AMPs.

In addition to engineering AMPs, sequence-activity mapping via SAMP-Dep provided mechanistic insight. Oncocin_1-15_ exhibits intracellular activity equivalent to full-length oncocin whereas oncocin_1-12_, oncocin_1-13_, and oncocin_1-14_ exhibit negligible activity. This provides additional insight to a previous C-terminal study^27^ that demonstrated sites 13-19 as useful for internalization yet not requisite for intracellular activity. Similarly, additional structural studies^26^ suggested a lack of ribosome engagement beyond site 13. Our data suggest that site 15 – and perhaps 14 and 13 – is requisite for intracellular activity; therefore, loss of extracellular activity was not decoupled from loss of intracellular activity *in vivo*.^27^

Quantifying a continuum of antimicrobial activity via SAMP-Dep empowers both engineering efforts and mechanistic understanding by deeply elucidating the AMP sequence-activity landscape. By providing a new high-throughput platform for engineering intracellularly-targeting AMPs, we believe SAMP-Dep is poised to help combat rising antimicrobial resistance. Success of SAMP-Dep potentiates deeply elucidating a new class of AMP sequence-activity relationships and guiding design of focused synthetic AMP libraries.

## Methods

### SAMP-Dep platform development

#### First-generation control synthesis

As controls, pETh vectors encoding oncocin (positive control) or a start followed by five stop codons (negative control), were synthesized. The first position of oncocin was mutated to methionine, ATG, to ensure translation. The oncocin sequence was codon optimized for *E. coli* with the IDT Codon Optimization Tool to aid efficient translation. To generate the pETh expression vectors encoding oncocin and negative control, pETh-Gp2-wt^42^ was double digested with NdeI (NEB) and BamHI-HF (NEB), agarose gel purified (Epoch Life Science), and assembled with oncocin and true negative oligo inserts (**Supplementary Table 2**) with NEBuilder^®^ HiFi DNA Assembly (NEB), and cloned in 5α *E. coli* cells (NEB). Constructs were validated by Sanger sequencing.

#### Induction test

50 μL of T7 Express LysY/I^q^ *E. coli* competent cells (NEB) were transformed with 80 ng of pETh oncocin or negative control, separately. After SOC outgrowth recovery, recovered cells were pelleted at 1,500 rcf for 3 minutes then suspended in 400 μL of lysogeny broth (LB) with 50 μg mL^-1^ kanamycin (kan). 50 μL of resuspended cells inoculated 4.95 mL of LB with 50 μg mL^-1^ kan and 0 to 0.50 mM isopropyl β-D-1 thiogalactopyranoside (IPTG). The liquid cultures were incubated at 37 °C with shaking at 250 rpm for nine hours. Optical density at 600 nm (OD600) was measured every 50 minutes with an Eppendorf Biophotometer plus.

#### Assay parameter determination

50 μL of T7 LysY/I^q^ *E. coli* competent cells (NEB) were transformed with 80 ng pETh oncocin or negative control, separately. During the SOC recovery, transformed cells were recovered in SOC with 0 or 0.50 mM IPTG. After recovery, recovered cells from each treatment were pelleted at 1,500 rcf for 3 minutes and suspended in 400 μL LB with 50 μg mL^-1^ kan. 50 μL of resuspended cells inoculated 4.95 mL of LB with 50 μg mL^-1^ kan with 0 or 0.50 mM IPTG prior to growth in a 30 or 37 °C incubator for 5 hours. After the outgrowth, 200 μL of each condition were added to a 96 well plate and incubated shaking at 282 rpm at 37 °C, and OD600 was monitored via a BioTek Synergy H1 microplate reader in 12-minute increments. Cell growth for each condition was monitored in six technical replicates.

### First-generation SAMP-Dep

#### First-generation library synthesis

To ensure adequate sequencing coverage, library size was tailored to Illumina HiSeq, which provided 220 million 2×50 paired end reads. The *n/n*+1 and *n/n*+2 first-generation library encoded (16 + 17) · (21^2^ - 1) = 14,520 unique amino acid mutants within (16 + 17) · (4^6^ - 1) = 135,135 DNA sequences, which provided ∼1,600 reads per DNA sequence per replicate per induction condition, assuming equal and full sequencing coverage.

To enable one-pot saturation mutagenesis, a BbvCI nicking endonuclease (NEB) recognition sequence was inserted into a non-coding region of the pETh oncocin plasmid to enable nicking mutagenesis (**Supplementary Table 2**).^43^ The pETh oncocin plasmid was digested with SphI (NEB), and the BbvCI recognition sequence was inserted via NEBuilder^®^ HiFi DNA Assembly (NEB).

Mutagenic primers for nicking mutagenesis were designed with Agilent QuikChange primer designer (**Supplementary Table 3**). The first-generation library was synthesized using nicking mutagenesis.^43^ The synthesized library was electroporated into MC1061 F- *E. coli* electrocompetent cells (Lucigen) then plated on a 245 × 245 mm LB and kan agar plate. After incubation overnight at 37 °C, 30 mL of LB with 50 μg mL^-1^ kan media suspended the plate culture. Plasmid DNA was extracted from 1 mL of suspended culture via miniprep (Epoch Life Science). The transformation yielded approximately 21 million colonies. The library was ∼155 times oversampled, ensuring coverage was not limited from transformation efficiency. The first-generation library was sequenced via Illumina MiSeq with v3 chemistry to quantify library composition. The parent oncocin sequence comprised 62% of the synthesized library, likely a result of remnant unmutated template and preferential amplification from wild-type primers.

#### First-generation SAMP-Dep protocol

To adequately sample the first-generation library, the transformation of T7 LysY/I^q^ *E. coli* (NEB) was scaled up to aid coverage. From the MiSeq library characterization, approximately 60% of the library was parental oncocin; therefore, assuming mutants other than parental were uniformly distributed, approximately 7.6 million transformed bacteria would provide 5.6-fold coverage of the first-generation library per induction condition per replicate, which would theoretically provide a >99% probability that every mutant would be observed at each induction condition per replicate. Given the T7 LysY/I^q^ *E. coli* transformation efficiency was approximately 560,000 CFU per 50 μL competent cells transformed with 80 ng pETh plasmid, sixteen 50 μL reactions of T7 LysY/I^q^ competent *E. coli* were transformed to yield a hypothetical 9 million CFU per replicate, providing adequate coverage of the first-generation library.

After recovery in SOC medium, cells were pelleted at 1,500 rcf and suspended in 400 μL LB with 50 μg mL^-1^ kan. 100 μL of the outgrowth suspension were added to 9.9 mL of LB with 50 μg mL^-1^ kan containing either 0, 0.15, 0.30, or 0.50 mM IPTG. Cells were incubated at 37 °C shaking at 250 rpm for 7.2 ± 0.1 hours. After incubation, cells were chilled on ice for ten minutes to slow growth, and the cells were pelleted by centrifuging at 3,000 rcf for five minutes. Supernatant was decanted, and cells from the same induction condition were combined. Cells for each condition were suspended in 5 mL of LB. DNA was extracted from 600 μL of each suspension via miniprep (Epoch Life Science). Three replicates were performed on three different days.

#### First-generation deep-sequencing sample preparation

To prepare the extracted plasmids for quantification via deep sequencing, 20 ng of extracted plasmid DNA was added to a 50 µL PCR reaction with Q5 DNA polymerase (NEB) for 16 cycles annealing at 53 °C. PCR primers were an equimolar mixture of FA and RA primers with 1N, 2N, or 3N offsets (**Supplementary Table 4**). After 16 cycles, 2 μL of 2 U L^-1^ ExoI (NEB) was added and incubated at 37 °C for 30 minutes then 80 °C for 20 minutes. A fresh 50 μL PCR reaction with Q5 (NEB) for 16 cycles annealing at 67 °C was mixed with 1 μL of ExoI (NEB) digested mixture as the template, FB primer, and a RB primer with a unique barcode indicating the induction condition and replicate number (**Supplementary Table 4**). The PCR product was agarose gel purified (Epoch Life Science) then deep sequenced via Illumina HiSeq 2500 with a high output 8-lane flow cell with v4 chemistry at the University of Minnesota Genomics Center.

#### First-generation deep sequence alignment and quality filtering

Computing resources from the Minnesota Supercomputing Institute were utilized with pandaseq^44^ to merge paired forward and reverse reads and trim primer sequences and with USearch^45^ to accept reads with less than 0.001 expected errors per sequence based on the quality score. To further filter for high quality reads, merged sequences were filtered for being observed at all uninduced conditions and had at least 100 read counts across all replicates and induction conditions. A total of 56 × 10^6^ reads (18 × 10^6^ from the first replicate,18 × 10^6^ from the second replicate, and 20 × 10^6^ from the third replicate) passed quality filtering.

#### Growth rate derivation

For cells growing exponentially, the growth rate constant of mutant i at IPTG concentration x, µ_i,x_, could be expressed as^46^,

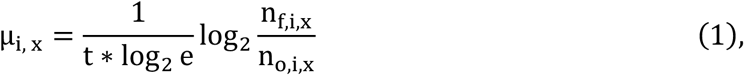

which was a function of t, the growth time; e, Euler’s number; n_f,i,x_, the final number of cells containing mutant i; and n_o,i,x_, the initial number cells containing mutant i at concentration x. Expanding the growth rate constant,

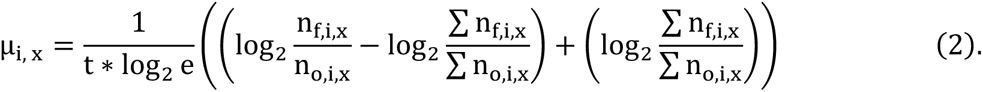

Within the expanded growth rate constant expression, the log_2_ enrichment of mutant i at IPTG concentration x, ε_i,x_, and bulk doubling times at IPTG concentration x, g_p,x_, were identified as

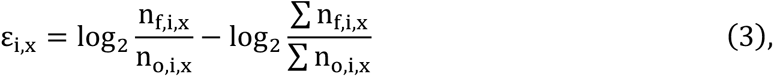

and

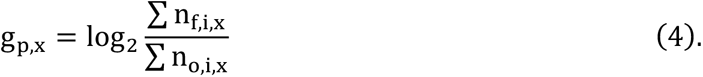

Substituting log_2_ enrichment and bulk doubling times into the expanded growth rate expression yielded

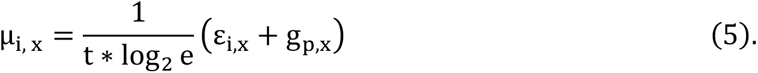

The log_2_ enrichment was immediately calculable; however, for the bulk number of doubling times at IPTG concentration x, several estimations were made. Null peptide growth rate constant at condition x was set equal to the null uninduced growth rate constant,

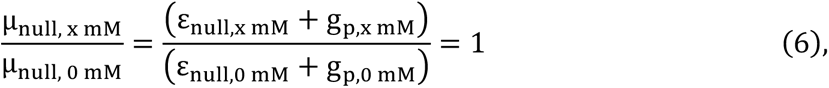

which was further simplified by setting ε_null,0 mM_to zero, for the negative control would not be enriched when not induced. g_p,0 mM_ was calculated from pilot experiments of uninduced T7 LysY/I^q^ growth (**Supplementary Figs. 1a and 3d**). Solving for g_p,x mM_, an expression for bulk doubling times was obtained

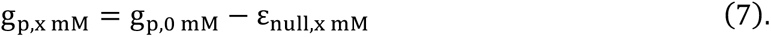

Substituting g_p,x mM_ into µ_i,x_ yielded

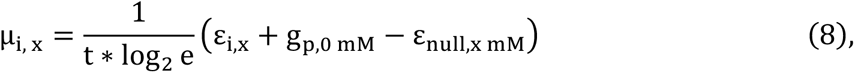

which gave the growth rate constant of the mutant i at induction level x, µ_i,x_, as a function of the growth time, t; log_2_ enrichment of mutant i at induction level x, ε_i,x_; the bulk number of doubling times at 0 mM IPTG, g_p,0 mM_; and the log_2_ enrichment of the negative control ε_null,x mM_

#### First-generation potency metric

For DNA based analysis, reads were tallied for unique DNA sequences, and for amino acid based analysis, reads were tallied for unique amino acid sequences. When summing counts per condition, a pseudo count of one was included for each induction condition to avoid log_2_(0) when calculating the log_2_ enrichment of the mutant abundance. The growth rate was calculated from Eq. (8). G_p,0 mM_ was approximated as 12.5 for an uninduced culture that grew 7.2 hours at a rate equal to uninduced oncocin in **Supplementary Figure 1a**. ε_null,x mM_ was the average log_2_ fold change of double stop mutants contained within oncocin positions 3-9 at inducer concentration X. The potency metric was the average growth rate:induction level slope. Mutations at M1 were not incorporated in library design and were likely deep sequencing errors, as evidenced by most M1 mutations having slope comparable to oncocin and all M1 mutations differing by only a single nucleotide (**Fig. 5a**).

### Second-generation SAMP-Dep

#### Second-generation library synthesis

The second-generation library diversity was codon optimized with SwiftLib^47^. Second-generation library inserts were synthesized by PCR using forward primers 1-16 and a common reverse primer (**Supplementary Table 5)**. Synthesized library inserts 1-16 were mixed in equimolar ratios and extended to the XbaI recognition site on the pET backbone using Q5 PCR (NEB) with the library insert as a template and XbaI extension and common reverse as primers (**Supplementary Table 5**). pETh-Gp2-wt^42^ was double digested with XbaI (NEB) and BamHI-HF (NEB), agarose gel purified (Epoch Life Science), ligated with T4 ligase (NEB) with the XbaI-extended mutagenic insert, purified with minElute (Qiagen), and electroporated into 5α *E. coli* electrocompetent cells (NEB). The transformation yielded approximately 2.1 million colonies; therefore, with a DNA diversity of 161,279 unique mutants, not including negative controls and oncocin, the library was ∼13 times oversampled, ensuring coverage was not limited by transformation efficiency. Negative control and oncocin inserts were ligated, transformed separately, and added to a final mole fraction of 0.01 each.

#### Second-generation SAMP-Dep protocol

The theoretical diversity of the second-generation library (161,664 DNA mutants, including negative controls and parental oncocin) was comparable to the first-generation (135,135 DNA mutants) and was expected to have more even distribution from the ligation synthesis; therefore, sixteen 50 μL reactions of T7 LysY/I^q^ competent *E. coli* competent cells (NEB) were transformed, providing adequate coverage.

The second-generation SAMP-Dep protocol followed the first-generation protocol with a few changes. First, transformed cells were induced at lower intermediate concentrations than the first-generation, 0, 0.10, 0.20, and 0.50 mM IPTG, to account for more producible second-generation parental transcript (**Supplementary Figs. 1a and 3d**). DNA was extracted after 6.7 ± 0.1 hours of growth to avoid saturation, for cell density accumulated quicker than the first-generation. After collecting cells from each induction condition and suspending in 5 mL LB, DNA from 400 μL of suspension was extracted via miniprep (Epoch Life Science). Three replicates were performed on three different days.

Samples were prepared as described for the first-generation library with modified FA and RA primers (**Supplementary Table 4**) and deep sequenced via Illumina NovaSeq with an SP flow cell at the University of Illinois Roy J. Carver Biotechnology Center.

#### Second-generation deep sequence alignment and quality filtering

Computing resources from the Minnesota Supercomputing Institute were utilized with USearch^45^ to merge forward and reverse reads, trim primer sequences, filter reads for less than 0.001 expected errors per sequence based on the quality score, find unique sequences, and match unique sequences with library design. To further filter for high quality reads, merged sequences were filtered for being observed at all uninduced conditions and had at least 40 read counts across all replicates and induction conditions. A total of 344 × 10^6^ reads (113 × 10^6^ from the first replicate, 103 × 10^6^ from the second replicate, and 128 × 10^6^ from the third replicate) passed quality filtering.

#### Second-generation potency metric analysis

A growth:induction slope was calculated as in the first-generation analysis, but to accurately quantify activity of potent mutants that were present when uninduced but were very depleted when slightly induced, the y-intercept was set to µ_i,0 mM_ when regressed. Moreover, to accurately quantify activity of highly active variants not observed when induced and to avoid log_2_(0) when calculating the log_2_ enrichment of the mutant abundance, a pseudo count of 10^-1^ was included for such cases. The pseudo count was 10^-1^ in attempt to quantify active mutants more accurately while not inflating slope error (**Supplementary Fig. 8a**). When extrapolating pseudo count for mutants with monotonically decreasing read count with induction and no observed read counts at full induction, a pseudo count of 10^-1^ was accurate for most mutants (**Supplementary Fig. 8b**). Although the true frequency varied for each mutant and would be quantifiable with deeper sequencing of induction conditions, a pseudo count of 10^-1^ matched clonal validation data well (**Supplementary Figs. 8c,d**). 10^-1^ correlated well with clonal data, indicating 10^-1^ did not underestimate the potency of variants unlike a pseudo count of 1, but clonal:SAMP-Dep slope was steep enough to indicate SAMP-Dep slope did not overestimate active mutant slope either unlike 10^-4^. Consistency between inter-replicate reproducibility and read count extrapolation indicated 10^-1^ accurately quantified active mutant activity. g _p,0 mM_ was approximated as 9.9 for an uninduced culture that grew 6.6 hours at a rate equal to uninduced oncocin in **Supplementary Figure 3d**. ε_null,x mM_ in Eq. 8 was the average log_2_ fold change of 384 negative control mutants (**Supplementary Table 5**). The potency metric was the average growth rate:induction level slope across replicates, as in the first-generation.

**Supplementary Figure 8.**
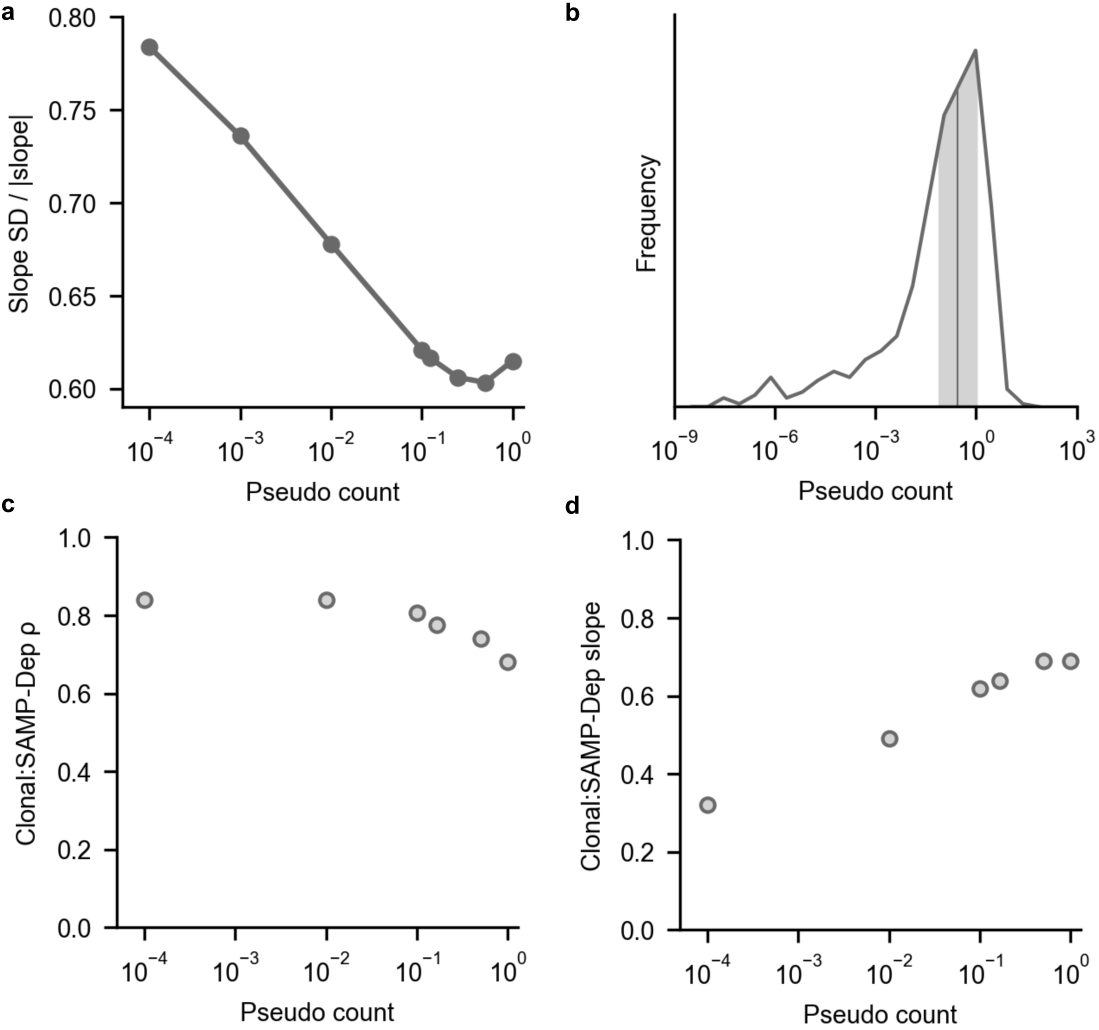
**A pseudo count of 10^-1^ accurately quantified potent mutant activity while maintaining low slope error. a**, Pseudo counts between 0.1 and 1 reduced relative error, average slope standard deviation (SD) relative to average slope, for mutants assigned a pseudo count. **b**, Mutants with monotonically decreasing read count with inducer concentration and not observed at full induction were extrapolated to predict read count at 0.50 mM IPTG. The median pseudo count was 0.3 (vertical line), and values ranging from 0.1 to 1 (shaded) were common. **c, d**, The relationship between clonally measured slope and SAMP-Dep slope (Fig. 4a) was calculated with different values for pseudo counts. **c,** Clonal and SAMP-Dep slopes correlated better with a low pseudo count, indicating high pseudo counts underestimated potent mutant activity. **d**, Clonal:SAMP-Dep slope ratio increased with larger pseudo counts, indicating SAMP-Dep slope was inaccurately large for low pseudo counts.

### SAMP-Dep validation

#### Clonal mutant synthesis

Individual mutant clones were selected for validation. Forward and reverse oligo pairs were ordered for each mutant with some overlaps in oligo sequence for both generations (**Supplementary Tables 6 and 7**). Inserts were assembled by PCR with Q5 polymerase (NEB), gel purified (Epoch Life Science), and inserted into the BamHI-HF (NEB) and NdeI (NEB) digested pETh backbone with NEBuilder^®^ HiFi DNA Assembly (NEB) and cloned in 5α *E. coli* cells (NEB).

#### Clonal SAMP-Dep

25 µL of T7 LysY/I^q^ *E. coli* competent cells (NEB) were transformed with 40 ng pETh mutant and recovered in 475 µL SOC. Cells were suspended in 1 mL LB and 50 µg mL^-1^ kan. 10 µL of suspension inoculated 190 µL of LB with 50 µg mL^-1^ kan with 0, 0.10, 0.20, 0.30, 0.40, or 0.50 mM IPTG in a 96 well plate. 200 µL water was added to perimeter wells to reduce evaporative losses. Absorbance was measured with a BioTek Synergy H1 microplate reader after incubation at 37 °C shaking at 250 rpm for 8.0 ± 0.5 hours. Growth rate was regressed from 0 mM to the lowest inducer concentration at which there was no quantifiable growth with a fixed y-intercept at µ_i,0 mM_ to calculate a clonal growth rate:induction slope. Clonal growth was measured at least in triplicate.

#### Minimal inhibitory concentration (MIC) assay

A single colony of T7 LysY/I^q^ *E. coli* (NEB) was suspended in 3 mL of LB and grown overnight. 10 µL of overnight culture inoculated 3 mL of Mueller Hinton Broth (MHB) (Sigma-Aldrich) and was grown at 37 °C shaking at 250 rpm for two hours. The outgrowth was diluted in MHB to a concentration of 5 × 10^5^ CFU mL^-1^. Peptides (chemically synthesized by Genscript, **Supplementary Table 6**) were suspended in DI H_2_O, and 10 µL of peptide was added to each well of a 96 well plate in a 2-fold dilution series for final concentrations of 80 to 2.5 µM peptide after dilution with 90 µL cell culture. 200 µL of water was added to perimeter cells to reduce evaporative losses. Absorbance was measured with a BioTek Synergy H1 microplate reader after incubation at 37 °C shaking at 250 rpm for 18 ± 0.5 hours. MIC was quantified as the concentration at which OD600 was less than 0.20 for trials at which cell density decreased monotonically with increasing peptide concentration. MIC was measured over at least six replicates.

To assess confidence in MIC values, a probability density function of the true MIC, µ, was approximated given a set of N experimental MIC values, X, as the marginal distribution of MIC independent of standard deviation of MIC, σ:

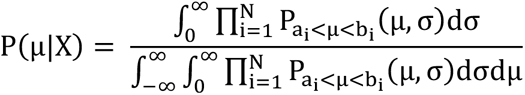

where P_a<μ<b_ was the probability that the true MIC, µ, was within the interval a_i_ and b_i_, such that a_i_ was the highest concentration of peptide to observe growth and b_i_ was the lowest concentration to ablate growth in an experimental MIC within X.^14^ Assuming the underlying distribution of noise was normally distributed around the mean,

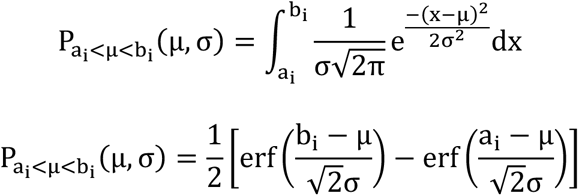

which provided a probability density distribution of MIC as a function of the experimental measurements.

The MIC was calculated as the mean of the distribution, and to assess significance of the mean MIC relative to oncocin, a bootstrapping approach performed 100,000 trials randomly sampling MIC values from the marginal distribution of the mutant and oncocin. The p value was calculated as the frequency of the resampled mutant MIC being greater than oncocin across the 100,000 trials.

### Sequence-activity relationships

#### Additive models for predicting SAMP-Dep slope

Amino acid sequences were represented by one-hot encoded vectors, paired with SAMP-Dep slope, and linearly regressed via LinearRegression from sklearn.linear_model. The linear coefficients were used to predict SAMP-Dep slope for new peptide sequences.

#### Slope interaction to quantify epistasis

To quantify interaction between mutations, the deviation from additivity was quantified between double and single mutants. The slope interaction was calculated as,

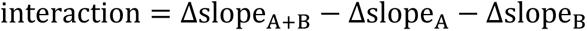

Where Δslope was the difference relative to parental oncocin. Therefore, negative interactions indicate the double mutant was more potent than expected from single mutants while positive interaction indicated a less potent double mutant than expected assuming additivity. Although growth:induction slope did not explicitly measure ΔG, we hypothesized an analogous analysis with slope could yield context-specific mutational effects. Slope had a limited dynamic range, for once ΔG_binding_ was unfavorable enough, cells would grow without detectable inhibition, plateauing slope at 0 M^-1^min^-1^. Additionally, independent mutational effects were not necessarily additive like ΔG_binding_.

#### Codon enrichment distribution and significance calculation

The enrichment of variants with comparable growth:induction slope for a given codon relative to all synonymous variants (*i.e.* DNA slope vs. amino acid slope in **Fig. 5d**) was approximated via bootstrapping via resampling with replacement for 5,000 trials. For each trial, a codon enrichment value was calculated as the frequency of codons present in non-superior DNA sequences (m_DNA_ > m_AA_ – 4 M^-1^min^-1^) relative to overall codon frequency. P value was the fraction of trials in which a codon enrichment value was either above or below one.

#### T-distributed stochastic neighbor embedding (t-SNE) of sequence space

To visualize higher-order second-generation sequence-space, a subset of mutants was visualized via t-SNE^37^. A group of 291 mutants were randomly sampled and placed in 5 M^-1^min^-1^ bins ranging from 5 to -40 M^-1^min-1 containing no more than 40 mutants to form an approximately uniform SAMP-Dep slope distribution. Mutants were one-hot encoded into vector representations of amino acid sequence, which was subsequently reduced to two dimensions via t-SNE. Clusters of comparable activity were identified across trajectories of t-SNE space (**Supplementary Fig. 6g**). Within each cluster, consensus mutations were defined as those present in greater than half of mutants in a cluster.

### Deep sequencing post hoc analysis

Precision and sensitivity quantified performance in experimental design. Genuine positives were defined as mutants that had p < 0.05 from a 1-tailed t-test relative to oncocin in the full data set, and genuine negatives had p ≥ 0.05. To assess the impact of replicates on SAMP-Dep robustness, replicates were removed, average slope and slope standard deviation recalculated, and a 1-tailed t-test performed relative to oncocin. Mutants that had p < 0.05 were predicted positive, and precision and sensitivity were calculated as the fraction of positives that were truly positive and the fraction of true positives that were positive, respectively. Similarly, for reducing induction conditions, slope, error, and significance were recalculated by regressing across a subset of induction concentrations, and precision and sensitivity for each subset was calculated. Resampling with less sequencing depth was modeled as,

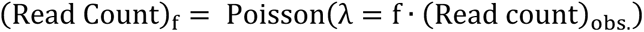

where f was the fraction of counts retained, and precision and sensitivity were calculated for each fraction.

## Acknowledgements

This work was supported by the National Institutes of Health (R01GM121777). We thank the University of Minnesota Genomics Center and the University of Illinois Roy J. Carver Biotechnology Center for assistance with deep sequencing. We thank the Minnesota Supercomputing Institute (MSI) for computational support. We also thank Dr. Pin-Kuang Lai for helpful discussions for oncocin sequence-function relationships and library design, Dr. Justin Klesmith for guidance with nicking mutagenesis, and Dr. Jorden Johnson for helpful discussions and feedback on the manuscript.

## Author Contributions

Conceptualization, M.P.D., S.C.R., and B.J.H.; Methodology, M.P.D., S.C.R., D.T.T., and B.J.H.; Software, M.P.D., S.C.R., K.A.F., and A.W.G.; Formal analysis, M.P.D. and A.W.G.; Investigation, M.P.D. and D.T.T.; Resources, M.P.D. and K.A.F.; Data curation, M.P.D.; Visualization, M.P.D.; Writing: original draft, M.P.D., B.J.H.; Writing: review & editing, M.P.D., S.C.R., K.A.F., D.T.T., A.W.G., B.J.H.; Funding acquisition, B.J.H.

