## Supplemental Information for "A platform for deep sequence-activity mapping and engineering antimicrobial peptides"

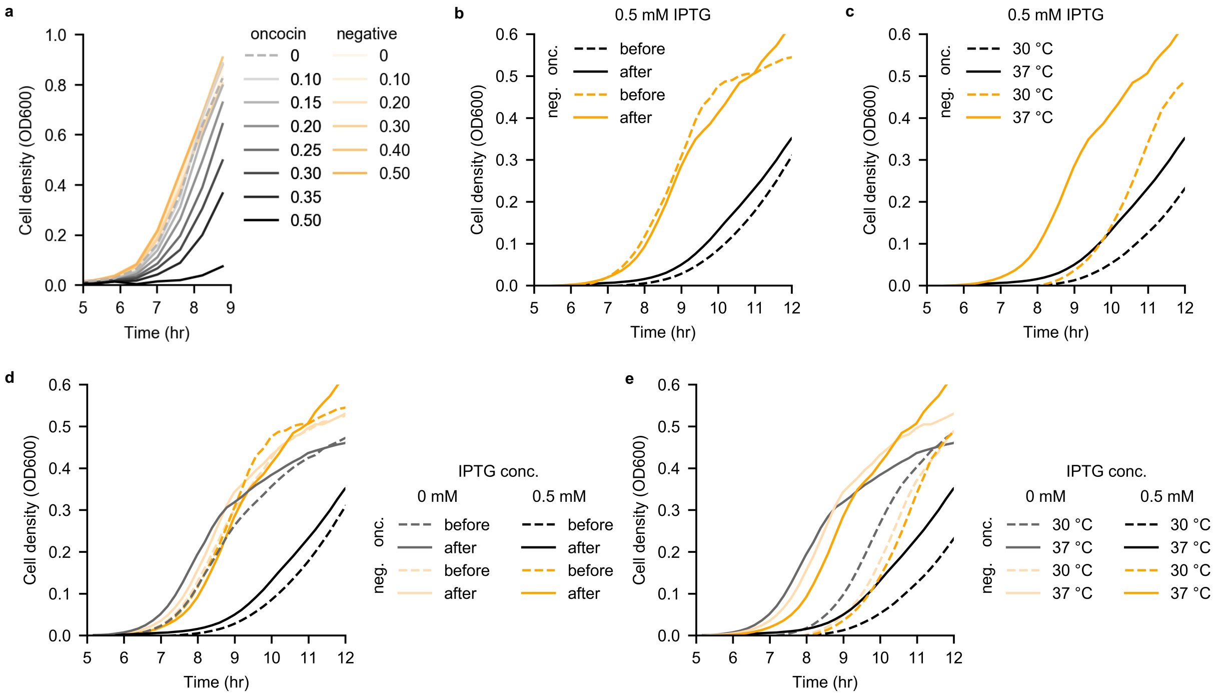


**Supplementary Figure 1. Assay development. a**, Expression of oncocin (black) and negative control (vector expressing a start codon followed by five stop codons; orange) in T7 LysY $I^q$ *E. coli* across varying induction levels revealed inducible depletion of oncocin. AMP production was induced at 37 ºC after transformation recovery. The legend indicates the concentration of IPTG inducer in mM. Uninduced oncocin was plotted with a dashed line to aid visualization. **b**, Induction start time did not substantially affect oncocin (black) depletion relative to negative control (orange). AMP production was induced with 0.50 mM IPTG before transformation recovery (dashed) or after recovery (solid). Cells were grown at 37 ºC. **c**, Lower induction temperature reduced depletion of oncocin (black) relative to negative control (orange). Cells were grown at 30 ºC (dashed) or 37 ºC (solid). AMP production was induced with 0.50 mM IPTG after the transformation recovery. **d**, **Supplementary Figure 1b** with uninduced populations (light). **e**, **Supplementary Figure 1c** with uninduced populations (light).

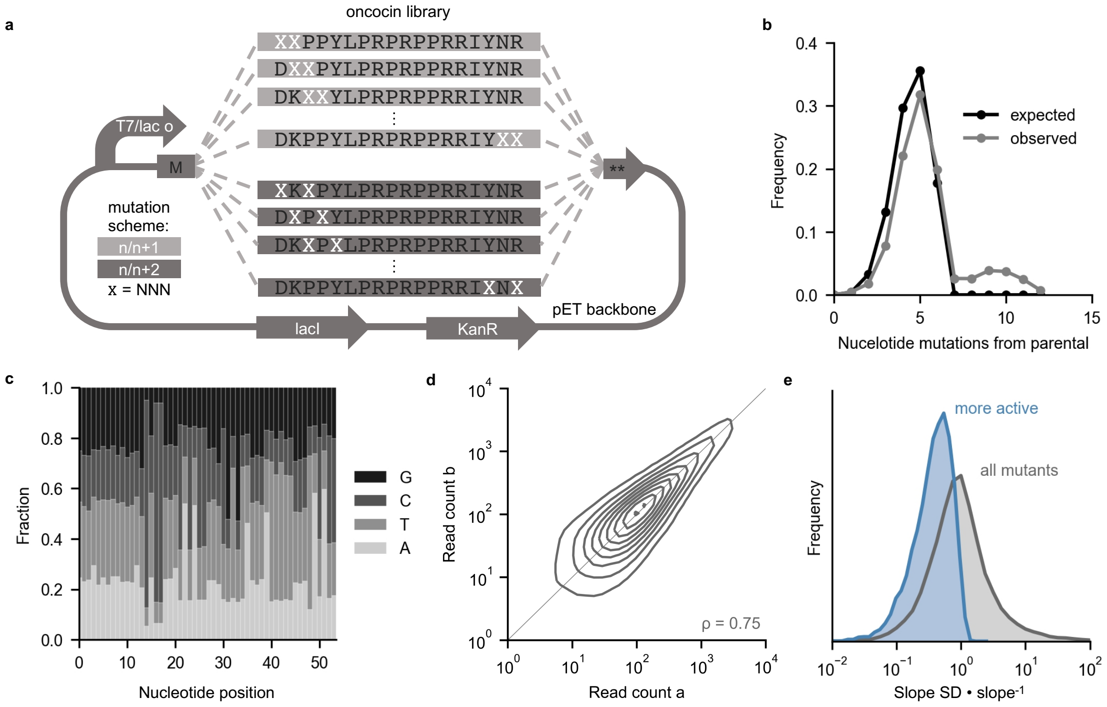


**Supplementary Figure 2. First-generation library design and deep sequencing analysis. a,** The first-generation oncocin mutagenic library used an *n*/*n*+1 and *n*/*n*+2 mutation scheme. Degenerate NNN codons (X) cover all 64 codons encoding 20 amino acids. **b**, The observed nucleotide mutation distribution followed the expected binomial distribution with $n_{mutations} = 6$ and $p_{mutation} = 0.75$. The number of nucleotide mutations was calculated relative to the parental oncocin DNA sequence. **c**, Mutated codons displayed the expected nucleotide composition from NNN codons. **d**, Read counts for each induction condition correlated across replicates. Read count a and b were randomly selected replicates without replacement for each mutant at each induction condition and represented as a kernel density plot. ρ indicates Pearson correlation coefficient. **e**, Slope was reproducible for most mutants more active than parental (blue). Relative slope standard deviation was higher for the full set of mutants due to many mutants having slope near 0 $M^{-1}min^{-1}$. Slope standard deviation (SD) was from three SAMP-Dep replicates.

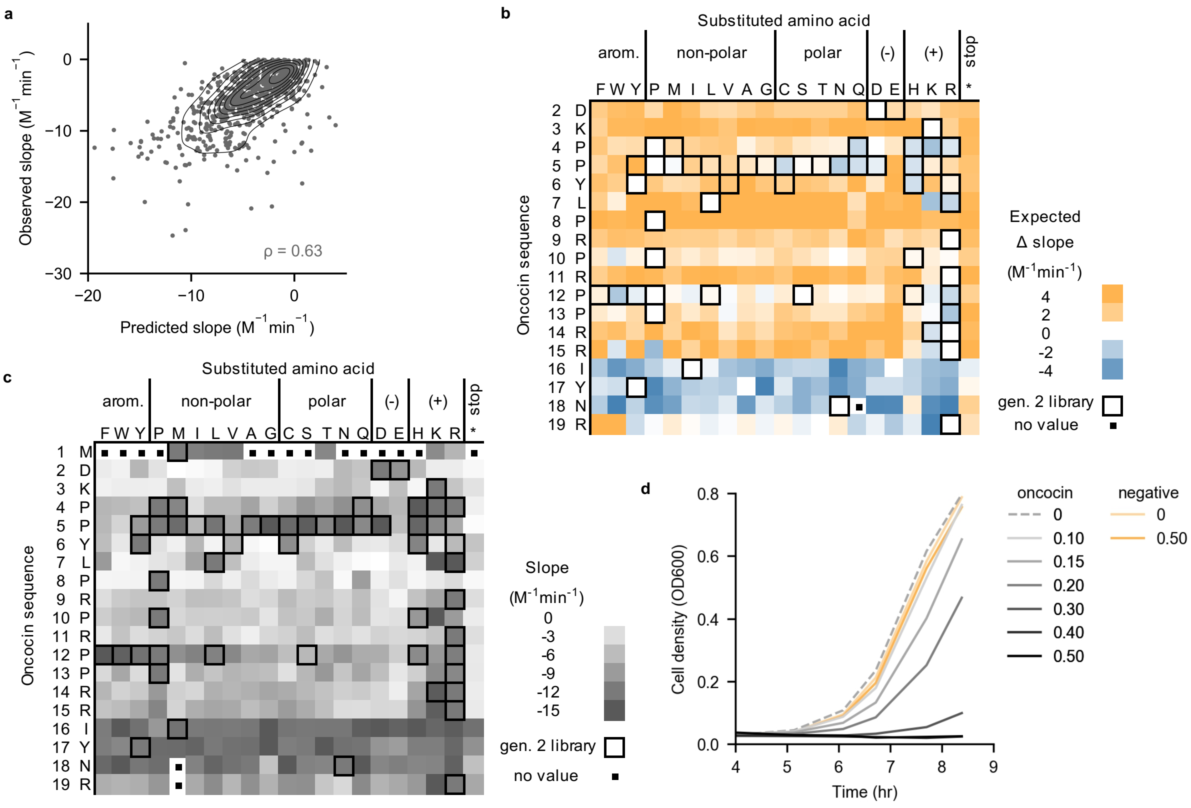


**Supplementary Figure 3. First-generation deep sequencing analysis empowers second-generation library design. a**, An additive model trained on 80% of first-generation sequence-activity pairs predicted slope for the remaining 20% of first-generation sequences. All mutants were filtered for slope standard deviation < 2 M$^{-1}$min$^{-1}$. **b**, Coefficients from an additive model trained on 100% of first-generation sequence-activity pairs indicated expected slope contributions. A negative expected change in slope (blue) suggested the mutation enhanced activity broadly. Outlined residues were incorporated in the second-generation library corresponding to **Figure 3a**. **c**, The single mutant slope heatmap indicated individual mutation effect on activity. Outlined residues were incorporated in the second-generation library corresponding to **Figure 3a**. **d**, Expression of second-generation parental oncocin (black) and negative control (orange) in T7 LysY I$^{q}$ *E. coli* across varying induction levels revealed inducible depletion of oncocin. AMP production was induced at 37 ºC after transformation recovery. The legend indicates the concentration of IPTG inducer in mM. Uninduced oncocin was plotted with a dashed line to aid visualization.

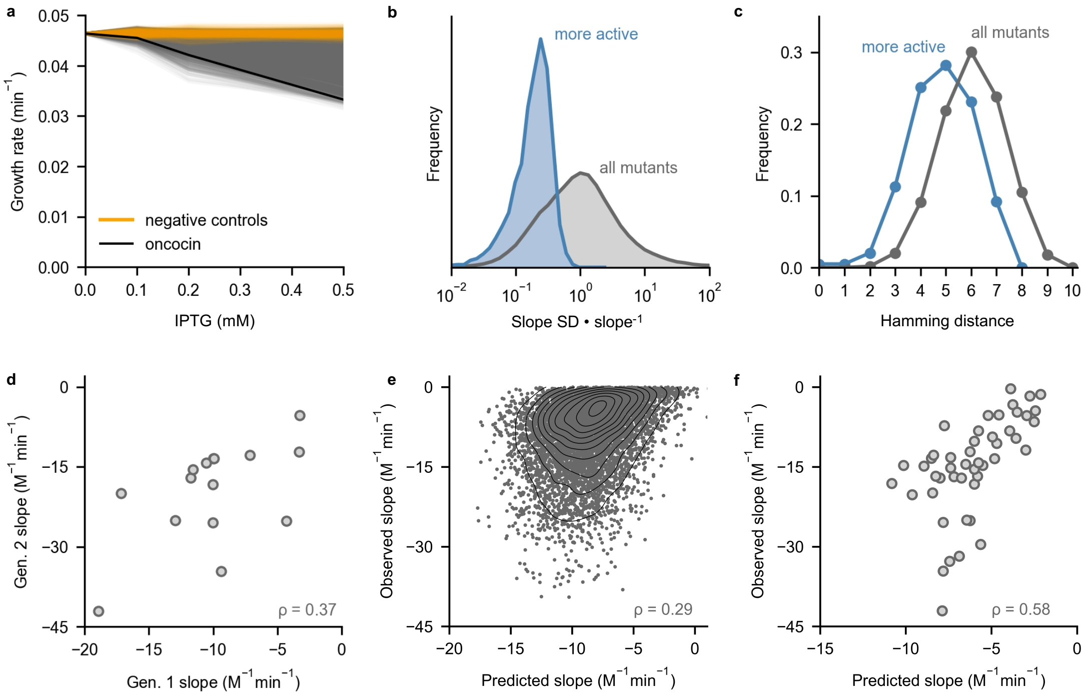


**Supplementary Figure 4. Second-generation deep sequencing analysis.** All mutants used for training and predictive analyses were filtered for slope standard deviation < 2 M$^{-1}$min$^{-1}$. **a**, Growth rate versus induction level for a subset of 10,000 mutants indicated most mutants reduced activity while some enhanced activity. Negative controls (orange) and parental oncocin (black) were included for comparison. All mutants had error comparable to parental oncocin (slope standard deviation < 2 M$^{-1}$min$^{-1}$). **b**, Slope was reproducible for most mutants more active than oncocin (blue). Slope reproducibility for all mutants (grey) was higher due to many mutants having slope near 0 M$^{-1}$min$^{-1}$. Slope standard deviation (SD) was from three SAMP-Dep replicates. **c**, More active mutants than oncocin (blue) were more sequence-similar to oncocin relative to the entire library (grey). Hamming distance was calculated on an amino acid basis relative to oncocin. **d**, Second-generation slope correlated with first-generation slope for amino acid variants observed in both libraries. **e**, An additive model trained on 100% of first-generation mutants predicted slope for 20% of second-generation sequences. The same subset of second-generation mutant slopes was predicted in **Figure 4d**. **f**, An additive model trained on 100% of first-generation mutants predicted slope of second-generation mutants that conformed to the first-generation library design.

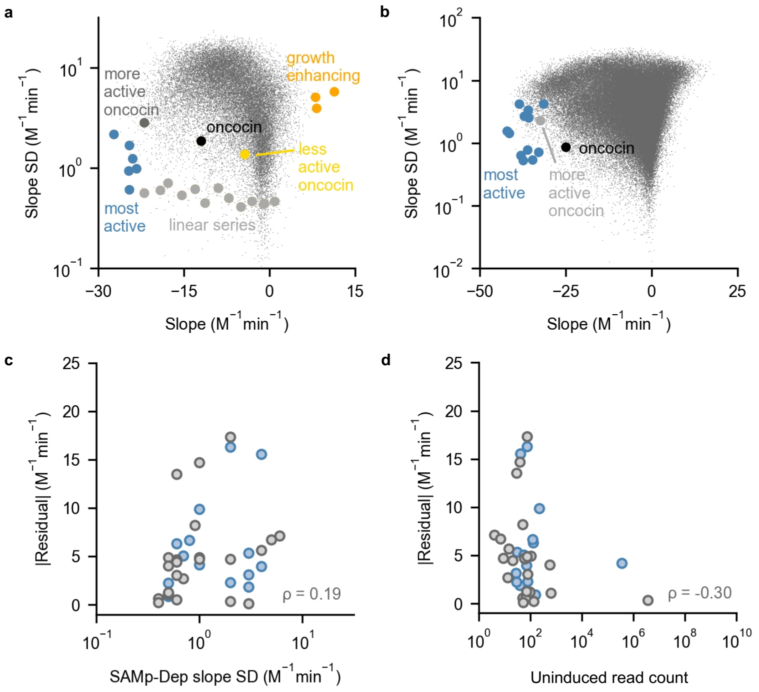


**Supplementary Figure 5. Clonal validation mutants and deviation from linear fit. a**, First-generation mutants selected for clonal validation answered several questions. The linear series (grey) addressed SAMP-Dep sensitivity and correlation with clonal activity. The most potent mutants (blue) addressed SAMP-Dep utility. Synonymous oncocin mutants (dark grey and yellow) addressed codon effect on activity. Putative growth enhancing mutants (orange) with positive SAMP-Dep slopes addressed an unexpected growth-enhancing phenotype. SD represents standard deviation. **b**, Second-generation mutants with substantially enhanced SAMP-Dep slope (blue) were selected for clonal validation. A synonymous oncocin mutant (grey) was among the most potent mutants. **c**, Residuals for all first-generation (grey) and second-generation (blue) clonal mutants relative to SAMP-Dep slope standard deviation indicated a weak relationship. Slope standard deviation was from three SAMP-Dep replicates. **d**, Residuals for all first-generation (grey) and second-generation (blue) clonal mutants relative to uninduced read count indicated a weak relationship.

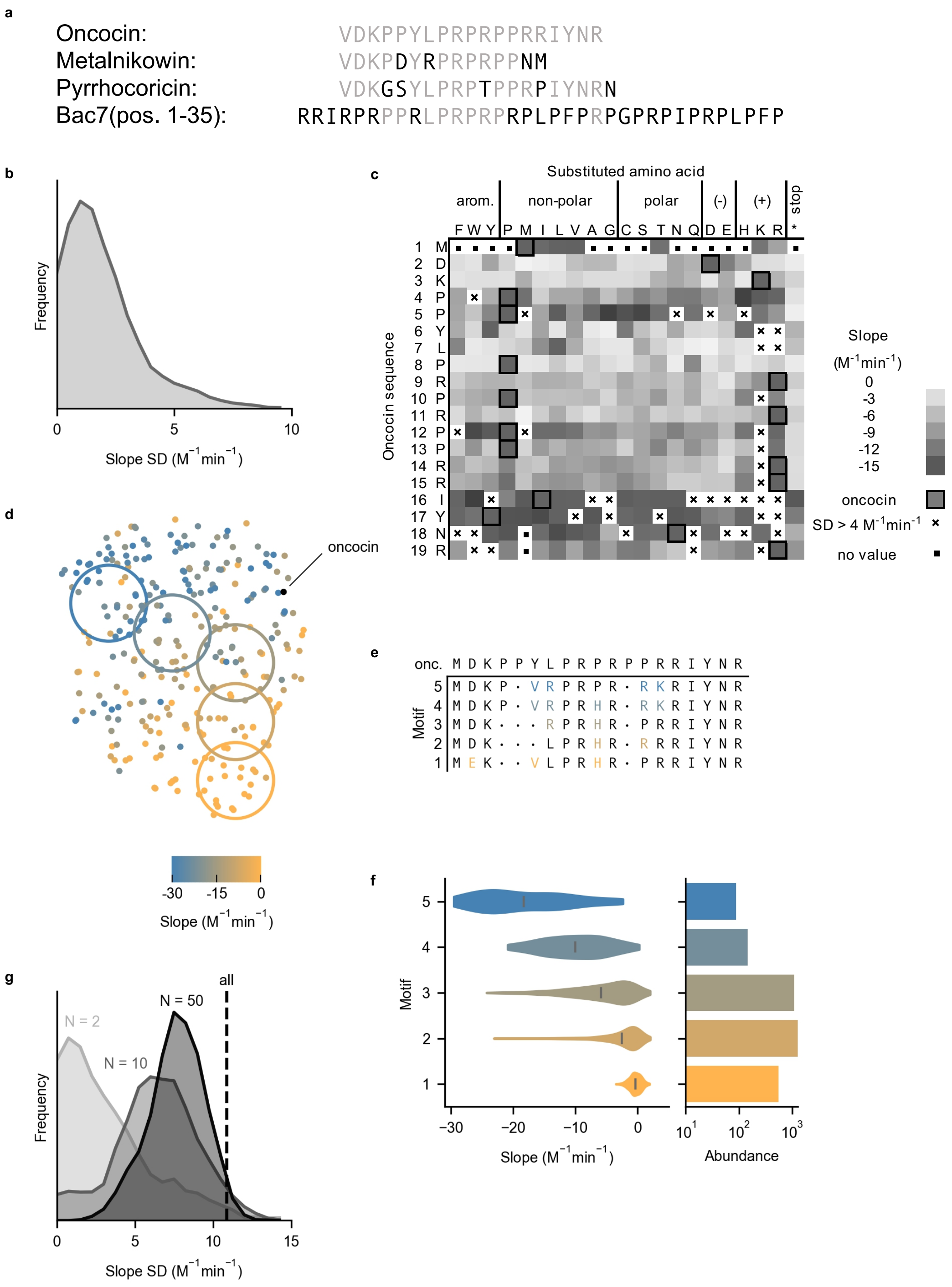


**Supplementary Figure 6. Additional first-generation sequence-function relationships. a**, Oncocin homologs metalnikowin, pyrrhocoricin, and bac7 have highly similar sequences. **b**, The growth:induction SAMP-Dep slope was reproducible for most mutants in the single mutation slope heatmap (**Fig. 5a**). The slope standard deviation from three SAMP-Dep replicates is presented with kernel smoothing. **c**, The slope of single-site mutants, from **Figure 5a**, with standard deviation < 4 $M^{-1}min^{-1}$. **d**, Alternative trajectory across a two-dimensional representation of sequence space (**Fig. 5f**). **e**, Common motifs conserved within colored bins from **Supplementary Figure 6d**. **f**, *Left*: Second-generation mutant slope distribution with motifs identified in **Supplementary Figure 6e**. *Right*: Motif prevalence in the second-generation library. **g**, Slope standard deviation histograms for N = 2, 10, and 50 nearest neighbors by Euclidean distance supported clustering. The dashed line represents slope standard deviation of all 291 mutants. The slope standard deviation is presented with kernel smoothing.

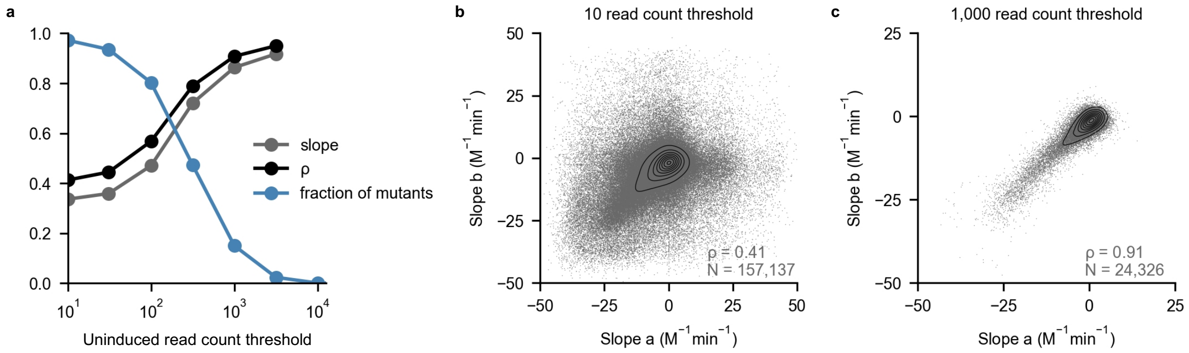


**Supplementary Figure 7. Slope reproducibility increased with mutant abundance.** **a**, Second-generation SAMP-Dep slope reproducibility across replicates increased with more stringent uninduced count thresholds. Slope (grey) and ρ (black) indicated the slope and Pearson correlation coefficient of SAMP-Dep slope between replicates. Fraction of mutants (blue) indicated the proportion that passed the uninduced read count threshold. **b**, Slope reproducibility was poor across replicates for mutants observed at least ten times when uninduced. **c**, Slope reproducibility was strong across replicates for mutants observed at least a thousand times when uninduced.

**Supplementary Table 1 | Validation results**

| DNA name* | Peptide name | Generation | Clonal slope ($M^{-1}min^{-1}$) | SAMP-Dep slope ($M^{-1}min^{-1}$) | MIC (µM) |
|---|---|---|---|---|---|
| clone 1 | A | 2 | -42 ± 2 | -42.0 ± 2 | 10 ± 1 |
| clone 2 | | 2 | -35 ± 5 | -41 ± 1 | |
| clone 3 | E | 2 | -30 ± 5 | -38.0 ± 0.6 | 22 ± 3 |
| clone 4 | | 2 | -28.8 ± 0.4 | -36.1 ± 0.8 | |
| clone 5 | | 2 | -27 ± 1 | -36 ± 3 | |
| clone 6 | | 2 | -25 ± 2 | -32.8 ± 0.7 | |
| clone 7 | | 2 | -24 ± 2 | -37.4 ± 0.5 | |
| clone 8 | D | 2 | -24 ± 2 | -36 ± 3 | 21 ± 4 |
| clone 9 | | 2 | -23.4 ± 0.4 | -34.5 ± 0.5 | |
| clone 10 | B | 2 | -23 ± 2 | -31 ± 4 | 15 ± 2 |
| clone 11 | | 2 | -22 ± 2 | -32 ± 2 | |
| clone 12 | C | 1 | -20 ± 2 | -24.5 ± 0.6 | 19 ± 4 |
| clone 13 | F | 2 | -20 ± 3 | -37 ± 3 | 60 ± 9 |
| parental onc. | oncocin | 2 | -19 ± 4 | -25 ± 1 | 25 ± 5 |
| clone 14 | | 1 | -16 ± 1 | -19.1 ± 0.6 | |
| clone 15 | | 1 | -13 ± 5 | -22 ± 3 | |
| clone 16 | | 1 | -10 ± 5 | -25 ± 2 | |
| clone 17 | | 1 | -9 ± 2 | -23 ± 1 | |
| clone 18 | | 1 | -8 ± 1 | -11.3 ± 0.5 | |
| parental onc. | oncocin | 1 | -8 ± 1 | -12 ± 2 | 25 ± 5 |
| clone 19 | | 2 | -8 ± 1 | -38 ± 4 | |
| clone 20 | | 1 | -8 ± 2 | -17.8 ± 0.7 | |
| clone 21 | | 1 | -7 ± 2 | -24.7 ± 0.9 | |
| clone 22 | | 1 | -7 ± 2 | -4 ± 1 | |
| clone 23 | | 1 | -5 ± 1 | -9.0 ± 0.6 | |
| clone 24 | | 1 | -4.8 ± 0.4 | -13.0 ± 0.6 | |
| clone 25 | | 1 | -4 ± 1 | -15.4 ± 0.5 | |
| clone 26 | | 1 | -2 ± 1 | -5.0 ± 0.4 | |
| clone 27 | | 1 | -1.5 ± 0.4 | 8 ± 5 | |
| clone 28 | | 1 | -1 ± 1 | -1.0 ± 0.4 | |
| clone 29 | | 1 | -1 ± 1 | -3.1 ± 0.5 | |
| clone 30 | G (negative) | 1 | -0.6 ± 0.4 | 0.8 ± 0.5 | >80 |
| clone 31 | | 1 | -0.2 ± 0.4 | -7.2 ± 0.5 | |
| clone 32 | | 1 | 0 ± 1 | 8 ± 4 | |
| clone 33 | | 1 | 0 ± 1 | -24 ± 1 | |
| clone 34 | | 1 | 0.6 ± 0.2 | -27 ± 2 | |
| negative | | 2 | 0.6 ± 0.2 | 0 ± 4 | |
| clone 35 | | 1 | 0.7 ± 0.2 | -21.0 ± 0.6 | |
| negative | | 1 | 0.7 ± 0.4 | -2 ± 2 | |
| clone 36 | | 1 | 1.0 ± 1 | 12 ± 6 | |

* DNA and peptide sequences in **Supplementary Table 6**

**Supplementary Table 2 | Key Resources.**

| Reagent | Source |
|---|---|
| Bacterial Strains | |
| *E. coli*, MC1061 F- | Lucigen |
| *E. coli*, T7 express LysY/$I^q$ | NEB |
| *E. coli*, 5-alpha | NEB |
| Critical commercial assays | |
| 300-bp Paired-end (2x300 PE) MiSeq with v3 chemistry | Illumina |
| 50-bp Paired-end (2x50 PE) HiSeq 2500 with v4 chemistry | Illumina |
| 150-bp Paired-end (2x150 PE) NovaSeq 6000 with SP flow cell | Illumina |
| Oligonucleotides | |
| pET oncocin V1M insert: TTAAGAAGGAGATATACATAATGGATAAGCCACCGTATTTACCACGTCCCCGCCCCCCACGTCGCATTTACAACCGCTGATAAGATCCCACCATCACCATCAT | IDT |
| pET true negative control insert: TTAAGAAGGAGATATACATAATGTGATAATGATAATGAGATCCCACCATCACCATCAT | IDT |
| BbvCI recognition site insert: GCCGCAAGGAATGGTGCATGCCTCAGCCATGCAAGGAGATGGCGCCC | IDT |
| First-generation one-pot saturation mutagenesis primers, see **Supplementary Table 3** | IDT |

| | |
|---|---|
| Illumina sequencing primers, see **Supplementary Table 4** | IDT |
| Second-generation library primers, see **Supplementary Table 5** | IDT |
| Clonal validation library primers, see **Supplementary Tables 6** and **7** | IDT |
| Peptides | |
| Peptide validation library, see **Supplementary Table 6** | Genscript |

**Supplementary Table 3 | First-generation one-pot saturation mutagenesis oligos**

| Degenerate codon positions | Sequence |
|---|---|
| 2/3 | ATAATTTTGTTTAACTTTAAGAAGGAGATATACATAATGNNNNNNCCAC CGTATTTACCACGTCCCCGCCCC |
| 3/4 | AGAAATAATTTTGTTTAACTTTAAGAAGGAGATATACATAATGGATNNN NNNCCGTATTTACCACGTCCCCGCCCC |
| 4/5 | TTTAACTTTAAGAAGGAGATATACATAATGGATAAGNNNNNNTATTTAC CACGTCCCCGCCCCCCACG |
| 5/6 | AAGAAGGAGATATACATAATGGATAAGCCANNNNNNTTACCACGTCCCC GCCCCCCACG |
| 6/7 | ATACATAATGGATAAGCCACCGNNNNNNCCACGTCCCCGCCCCCCACGT C |
| 7/8 | AAGGAGATATACATAATGGATAAGCCACCGTATNNNNNNCGTCCCCGCC CCCCACG |
| 8/9 | AGGAGATATACATAATGGATAAGCCACCGTATTTANNNNNNCCCCGCCC CCCACGTCGC |
| 9/10 | GGAGATATACATAATGGATAAGCCACCGTATTTACCANNNNNNCGCCCC CCACGTCGCATTT |
| 10/11 | GAGATATACATAATGGATAAGCCACCGTATTTACCACGTNNNNNNCCCC CACGTCGCATTTACAA |
| 11/12 | GATAAGCCACCGTATTTACCACGTCCCNNNNNNCCACGTCGCATTTACA ACCGCTGATAA |

| | |
|---|---|
| 12/13 | GCCACCGTATTTACCACGTCCCCGCNNNNNNCGTCGCATTTACAACCGCTGATAAG |
| 13/14 | ACCGTATTTACCACGTCCCCGCCCCNNNNNNCGCATTTACAACCGCTGATAAGATC |
| 14/15 | CGTATTTACCACGTCCCCGCCCCCCANNNNNNATTTACAACCGCTGATAAGATCCCAC |
| 15/16 | CCACGTCCCCGCCCCCCACGTNNNNNNTACAACCGCTGATAAGATCCC |
| 16/17 | CGTCCCCGCCCCCCACGTCGCNNNNNNAACCGCTGATAAGATCCCACC |
| 17/18 | ACGTCCCCGCCCCCCACGTCGCATTNNNNNNCGCTGATAAGATCCCAC |
| 18/19 | CCCCGCCCCCCACGTCGCATTTACNNNNNNTGATAAGATCCCACCATCACCATC |
| 2/4 | TTTGTTTAACTTTAAGAAGGAGATATACATAATGNNNAAGNNNCCGTATTTACCACGTCCCCGCCCCCCA |
| 3/5 | GTTTAACTTTAAGAAGGAGATATACATAATGGATNNNCCANNNTATTTACCACGTCCCCGCCCCCCAC |
| 4/6 | CTTTAAGAAGGAGATATACATAATGGATAAGNNNCCGNNNTTACCACGTCCCCGCCCCCCAC |
| 5/7 | TTAACTTTAAGAAGGAGATATACATAATGGATAAGCCANNNTATNNNCCACGTCCCCGCCCCCCACG |
| 6/8 | GATATACATAATGGATAAGCCACCGNNNTTANNNCGTCCCCGCCCCCCACGTCGCATTT |
| 7/9 | GGAGATATACATAATGGATAAGCCACCGTATNNNCCANNNCCCCGCCCCCCACGTC |

| | |
|---|---|
| 8/10 | GAAGGAGATATACATAATGGATAAGCCACCGTATTTANNNCGTNNNCGCCCCCCACGTCGCATTT |
| 9/11 | GGAGATATACATAATGGATAAGCCACCGTATTTACCANNNCCCNNNCCCCCACGTCGCATTTACA |
| 10/12 | AATGGATAAGCCACCGTATTTACCACGTNNNCGCNNNCCACGTCGCATTTACAACCGCTG |
| 11/13 | TAAGCCACCGTATTTACCACGTCCCNNNCCCNNNCGTCGCATTTACAACCGCTGATAAG |
| 12/14 | GCCACCGTATTTACCACGTCCCCGCNNNCCANNNCGCATTTACAACCGCTGATAAGATC |
| 13/15 | CCGTATTTACCACGTCCCCGCCCCNNNCGTNNNATTTACAACCGCTGATAAGATCCC |
| 14/16 | TTACCACGTCCCCGCCCCCCANNNCGCNNNTACAACCGCTGATAAGATCCC |
| 15/17 | CCACGTCCCCGCCCCCCACGTNNNATTNNNAACCGCTGATAAGATCCCACC |
| 16/18 | CGTCCCCGCCCCCCACGTCGCNNNTACNNNCGCTGATAAGATCCCACCATC |
| 17/19 | ACGTCCCCGCCCCCCACGTCGCATTNNNAACNNNTGATAAGATCCCACCATCAC |

**Supplementary Table 4 | Illumina sequencing primers**

| Name | Sequence |
|---|---|
| $FA_{G1}$ | TTTCCCTACACGACGCTCTTCCGATCT[1 to 3 N] AGAAGGAGATATACATAATG |
| $RA_{G1}$ | GTTCAGACGTGTGCTCTTCCGATCT[1 to 3 N]GGTGGGATCTTATCA |
| $FA_{G2}$ | TTTCCCTACACGACGCTCTTCCGATCT[4 to 8 N] AGAAGGAGATATACATATGGA |
| $RA_{G2}$ | GTTCAGACGTGTGCTCTTCCGATCT[4 to 8 N]TGGTGGGATCCTCAT |
| FB | AATGATACGGCGACCACCGAGATCTACACTCTTTCCCTACACGACGCTCTT |
| RB | CAAGCAGAAGACGGCATACGAGAT[Barcode]GTGACTGGAGTTCAGACG TGTGCTCTTCC |

**Supplementary Table 5 | Second-generation library primers**

| Primer name | Sequence |
|---|---|
| $Fw1_{G2}$ | ATATACATATGGAMAAAADGKVCYRCCKTCCGCGTCMTCGTYSGC SAARACGTATTTACAACCGTTAATGAGGATCCCACCA |
| $Fw2_{G2}$ | ATATACATATGGAMAAAADGKVCYRCCKTCCGCGTCMTCGTYWCC SAARACGTATTTACAACCGTTAATGAGGATCCCACCA |
| $Fw3_{G2}$ | ATATACATATGGAMAAAADGMHSYRCCKTCCGCGTCMTCGTYSGC SAARACGTATTTACAACCGTTAATGAGGATCCCACCA |
| $Fw4_{G2}$ | ATATACATATGGAMAAACMMKVCYRCCKTCCGCGTCMTCGTYSGC SAARACGTATTTACAACCGTTAATGAGGATCCCACCA |

| | |
|---|---|
| Fw5$_{G2}$ | ATATACATATGGAMAAACMMMHSYRCCKTCCGCGTCMTCGTYSGC SAARACGTATTTACAACCGTTAATGAGGATCCCACCA |
| Fw6$_{G2}$ | ATATACATATGGAMAAACMMKVCYRCCKTCCGCGTCMTCGTYWCC SAARACGTATTTACAACCGTTAATGAGGATCCCACCA |
| Fw7$_{G2}$ | ATATACATATGGAMAAAADGMHSYRCCKTCCGCGTCMTCGTYWCC SAARACGTATTTACAACCGTTAATGAGGATCCCACCA |
| Fw8$_{G2}$ | ATATACATATGGAMAAACMMMHSYRCCKTCCGCGTCMTCGTYWCC SAARACGTATTTACAACCGTTAATGAGGATCCCACCA |
| Fw9$_{G2}$ | ATATACATATGGAMAAAADGKVCGTGCKTCCGCGTCMTCGTYSGC SAARACGTATTTACAACCGTTAATGAGGATCCCACCA |
| Fw10$_{G2}$ | ATATACATATGGAMAAAADGKVCGTGCKTCCGCGTCMTCGTYWCC SAARACGTATTTACAACCGTTAATGAGGATCCCACCA |
| Fw11$_{G2}$ | ATATACATATGGAMAAAADGMHSGTGCKTCCGCGTCMTCGTYSGC SAARACGTATTTACAACCGTTAATGAGGATCCCACCA |
| Fw12$_{G2}$ | ATATACATATGGAMAAACMMKVCGTGCKTCCGCGTCMTCGTYSGC SAARACGTATTTACAACCGTTAATGAGGATCCCACCA |
| Fw13$_{G2}$ | ATATACATATGGAMAAACMMMHSGTGCKTCCGCGTCMTCGTYSGC SAARACGTATTTACAACCGTTAATGAGGATCCCACCA |
| Fw14$_{G2}$ | ATATACATATGGAMAAACMMKVCGTGCKTCCGCGTCMTCGTYWCC SAARACGTATTTACAACCGTTAATGAGGATCCCACCA |
| Fw15$_{G2}$ | ATATACATATGGAMAAAADGMHSGTGCKTCCGCGTCMTCGTYWCC SAARACGTATTTACAACCGTTAATGAGGATCCCACCA |
| Fw16$_{G2}$ | ATATACATATGGAMAAACMMMHSGTGCKTCCGCGTCMTCGTYWCC SAARACGTATTTACAACCGTTAATGAGGATCCCACCA |

| | |
|---|---|
| Fw negative$_{G2}$ | ATATACATATGGAMTAAADGTAGYRCTAGCCGTAACMTTAGYSGT AGARATAAATTTAGAACTAGTAATGAGGATCCCACCA |
| Fw oncocin$_{G2}$ | ATATACATATGGACAAACCACCGTACCTTCCGCGTCCTCGTCCGC CAAGACGTATTTACAACCGTTAATGAGGATCCCACCA |
| Common Rv$_{G2}$ | TGGTGGGATCCTCATTAACGGTTGTAAATAC |
| XbaI extension | TTCCCCTCTAGAAATAATTTTGTTTAACTTTAAGAAGGAGATATA CATATGGAMAAAC |

**Supplementary Table 6 | Clonal validation library sequences and primers**

| DNA name* | Peptide name | Generation | DNA sequence | Peptide sequence | Forward primer* | Reverse primer* |
|---|---|---|---|---|---|---|
| clone 1 | A | 2 | ATGGACAAACCAAACCGCCTTCCGCGTCCTCGTCCGCCAAGACGTATTTACAACCGT | MDKPNRLPRPRPPRRIYNR | 2fw4 | 2rv1 |
| clone 2 | | 2 | ATGGACAAAAGGCCGGTGCGTCCGCGTCCTCGTTTCCCAAGACGTATTTACAACCGT | MDKRPVRPRPRFPRRIYNR | 2fw14 | 2rv11 |
| clone 3 | E | 2 | ATGGACAAACCCTCCTACCTTCCGCGTCCTCGTCTCCCAAGACGTATTTACAACCGT | MDKPSYLPRPRLPRRIYNR | 2fw11 | 2rv9 |
| clone 4 | | 2 | ATGGACAAACCACCGTGCCGTCCGCGTCCTCGTTGGCCAAGACGTATTTACAACCGT | MDKPPCRPRPRWPRRIYNR | 2fw10 | 2rv8 |
| clone 5 | | 2 | ATGGACAAAAGGCCCTACCGTCCGCGTCCTCGTCCGCCAAAACGTATTTACAACCGT | MDKRPYRPRPRPPKRIYNR | 2fw6 | 2rv4 |
| clone 6 | | 2 | ATGGACAAACCCCCGTACCGTCCGCGTCCTCGTTGGCGAAGACGTATTTACAACCGT | MDKPPYRPRPRWRRRIYNR | 2fw12 | 2rv10 |
| clone 7 | | 2 | ATGGACAAACCATGCCACCGTCCGCGTCCTCGTTACCGAAGACGTATTTACAACCGT | MDKPCHRPRPRYRRRIYNR | 2fw9 | 2rv7 |
| clone 8 | D | 2 | ATGGACAAACCACCCTACCGTCCGCGTCCTCGTTACCGAAAACGTATTTACAACCGT | MDKPPYRPRPRYRKRIYNR | 2fw5 | 2rv3 |
| clone 9 | | 2 | ATGGACAAAAGGCCCTACCTTCCGCGTCCTCGTTTCCGAAAACGTATTTACAACCGT | MDKRPYLPRPRFRKRIYNR | 2fw8 | 2rv6 |
| clone 10 | B | 2 | ATGGACAAACACCCGCACCGTCCGCGTCCTCGTTACCGAAGACGTATTTACAACCGT | MDKHPHRPRPRYRRRIYNR | 2fw13 | 2rv7 |
| clone 11 | | 2 | ATGGACAAACCCCCGTACCTTCCGCGTCCTCGTCCGCCAAGACGTATTTACAACCGT | MDKPPYLPRPRPPRRIYNR | 2fw2 | 2rv1 |
| clone 12 | C | 1 | ATGGATAAGCCACCGTATTTACCACGTCCCCGCCCCCCACGTCGCATTGGGAACAGA | MDKPPYLPRPRPPRRIGNR | 1fw19 | 1rv11 |
| clone 13 | F | 2 | ATGGACAAACCAACCCGCCTTCCGCGTCCTCGTCGGCCAAAACGTATTTACAACCGT | MDKPTRLPRPRRPKRIYNR | 2fw3 | 2rv2 |
| parental onc. | oncocin | 2 | ATGGACAAACCACCGTACCTTCCGCGTCCTCGTCCGCCAAGACGTATTTACAACCGT | MDKPPYLPRPRPPRRIYNR | 2fw1 | 2rv1 |
| clone 14 | | 1 | ATGGATAAGCCACCGTATTTACCACGTCCCCGCCCCCCACGGAGGATTTACAACCGC | MDKPPYLPRPRPPRRIYNR | 1fw11 | 1rv6 |
| clone 15 | | 1 | ATGGATAAGCCACCGTATTTACCACGGCCCCGGCCCCCACGTCGCATTTACAACCGC | MDKPPYLPRPRPPRRIYNR | 1fw17 | 1rv1 |
| clone 16 | | 1 | ATGGATAAGCCACCGTATTTACCACGTCCCCGCAGACGTCGTCGCATTTACAACCGC | MDKPPYLPRPRRRRRIYNR | 1fw24 | 1rv4 |
| clone 17 | | 1 | ATGGATAAGCCACCGTATTTACCACGTCCCCGCAATAGGCGTCGCATTTACAACCGC | MDKPPYLPRPRNRRRIYNR | 1fw22 | 1rv8 |
| clone 18 | | 1 | ATGGATAAGCCACCGTATTTACCACGTCCCCGCCCGCCACGTCGCATTTACAACCGC | MDKPPYLPRPRPPRRIYNR | 1fw7 | 1rv1 |
| parental onc. | oncocin | 1 | ATGGATAAGCCACCGTATTTACCACGTCCCCGCCCCCCACGTCGCATTTACAACCGC | MDKPPYLPRPRPPRRIYNR | 1fw16 | 1rv1 |
| clone 19 | | 2 | ATGGAAAAACCCACCCGCCTTCCGCGTCCTCGTCACCGAAAACGTATTTACAACCGT | MEKPTRLPRPRHRKRIYNR | 2fw7 | 2rv5 |
| clone 20 | | 1 | ATGGATAAGCCACCGTATTTACCACGTCCCCGCCCCCCACGTCGCCCTCGTAACCGC | MDKPPYLPRPRPPRRPRNR | 1fw10 | 1rv5 |
| clone 21 | | 1 | ATGGATAAGCCACCGTATTTACCACGTCCCCGCCGGGTGCGTCGCATTTACAACCGC | MDKPPYLPRPRRVRRIYNR | 1fw20 | 1rv8 |
| clone 22 | | 1 | ATGGATAAGCCACCGTATTTACCACGTCCCCGCCCTCCAAGACGCATTTACAACCGC | MDKPPYLPRPRPPRRIYNR | 1fw18 | 1rv10 |
| clone 23 | | 1 | ATGGATAAGCCACCGTATTTACCACGTCCCCGCCCCCCACGTCGCATTTACACCCGC | MDKPPYLPRPRPPRRIYTR | 1fw6 | 1rv3 |
| clone 24 | | 1 | ATGGATAAGCCACCGTATTTACCACGTCCCCGCTTTGTTCGTCGCATTTACAACCGC | MDKPPYLPRPRFVRRIYNR | 1fw8 | 1rv4 |
| clone 25 | | 1 | ATGGATAAGCCACCGTATTTACCACGTCCCCGCATGAAACGTCGCATTTACAACCGC | MDKPPYLPRPRMKRRIYNR | 1fw9 | 1rv1 |
| clone 26 | | 1 | ATGGATAAGCCACCGTATTTACCACGTCCCCGCTTACCAGGGCGCATTTACAACCGC | MDKPPYLPRPRLPGRIYNR | 1fw4 | 1rv2 |
| clone 27 | | 1 | ATGGATAAGCCACCGTATTTACCACGTCCCCGCCCCCCATTATGGATTTACAACCGC | MDKPPYLPRPRPPLWIYNR | 1fw15 | 1rv9 |
| clone 28 | | 1 | ATGGAGGTTCCACCGTATTTACCACGTCCCCGCCCCCCACGTCGCATTTACAACCGC | MEVPPYLPRPRPPRRIYNR | 1fw2 | 1rv1 |
| clone 29 | | 1 | ATGGATAAGCCACCGTATTTACCATCACCCATTCCCCCACGTCGCATTTACAACCGC | MDKPPYLPSPIPPRRIYNR | 1fw3 | 1rv1 |
| clone 30 | G (negative) | 1 | ATGGGCGAGCCACCGTATTTACCACGTCCCCGCCCCCCACGTCGCATTTACAACCGC | MGEPPYLPRPRPPRRIYNR | 1fw1 | 1rv1 |
| clone 31 | | 1 | ATGGATAAGCCACCGTATTTTCCACGTCCCCGCCCCCCACGTCGCATTTACAACCGC | MDKPPYFPRPRPPRRIYNR | 1fw5 | 1rv1 |
| clone 32 | | 1 | ATGGATAAGCCACCGTATTTACCACGTCCCGTGCCCGTGCGTCGCATTTACAACCGC | MDKPPYLPRPVPVRRIYNR | 1fw14 | 1rv8 |
| clone 33 | | 1 | ATGGATAAGCCACCGTATTTACCACGTACAATACCCCCACGTCGCATTTACAACCGC | MDKPPYLPRTIPPRRIYNR | 1fw21 | 1rv1 |
| clone 34 | | 1 | ATGGATAAGCCACCGTATGGTCCAGCTCCCCGCCCCCCATTGTGGATTTACAACCGC | MDKPPYGPAPRPPLWIYNR | 1fw23 | 1rv12 |
| negative | | 2 | ATGGAMTAGADGTAGYRCTAGCCGTAGCMTTAGYSGTAGARATAGATTTAGAACTAG | M• | see table _ | see table _ |
| clone 35 | | 1 | ATGGATAAGCCATGTTATCTTCCACGTCCCCGCCCCCCAGTATGGATTTACAACCGC | MDKPCYLPRPRPPVWIYNR | 1fw12 | 1rv7 |
| negative | | 1 | ATGTGATAATGATAATGA | M | see table _ | see table _ |
| clone 36 | | 1 | ATGGATAAGCCACCGTATTTACCAGTTTTCCGCCCCCCACGTCGCATTTACAACCGC | MDKPPYLPVFRPPRRIYNR | 1fw13 | 1rv1 |

* Primer sequences in **Supplementary Table 7**

**Supplementary Table 7 | Clonal validation library primers**

| Primer name | Sequence |
|---|---|
| 1fw1 | AAGGAGATATACATAATGGGCGAGCCACCGTATTTACCACGTCCCCGCCCCCCACGTCGCATTTACAACC |
| 1fw2 | AAGGAGATATACATAATGGAGGTTCCACCGTATTTACCACGTCCCCGCCCCCCACGTCGCATTTACAACC |
| 1fw3 | AAGGAGATATACATAATGGATAAGCCACCGTATTTACCATCACCCATTCCCCCACGTCGCATTTACAACC |
| 1fw4 | AAGGAGATATACATAATGGATAAGCCACCGTATTTACCACGTCCCCGCTTACCAGGGCGCATTTACAACC |
| 1fw5 | AAGGAGATATACATAATGGATAAGCCACCGTATTTTCCACGTCCCCGCCCCCCACGTCGCATTTACAACC |
| 1fw6 | AAGGAGATATACATAATGGATAAGCCACCGTATTTACCACGTCCCCGCCCCCCACGTCGCATTTACACCC |
| 1fw7 | AAGGAGATATACATAATGGATAAGCCACCGTATTTACCACGTCCCCGCCCGCCACGTCGCATTTACAACC |
| 1fw8 | AAGGAGATATACATAATGGATAAGCCACCGTATTTACCACGTCCCCGCTTTGTTCGTCGCATTTACAACC |
| 1fw9 | AAGGAGATATACATAATGGATAAGCCACCGTATTTACCACGTCCCCGCATGAAACGTCGCATTTACAACC |
| 1fw10 | AAGGAGATATACATAATGGATAAGCCACCGTATTTACCACGTCCCCGCCCCCCACGTCGCCCTCGTAACC |
| 1fw11 | AAGGAGATATACATAATGGATAAGCCACCGTATTTACCACGTCCCCGCCCCCCACGGAGGATTTACAACC |
| 1fw12 | AAGGAGATATACATAATGGATAAGCCATGTTATCTTCCACGTCCCCGCCCCCCAGTATGGATTTACAACCGCTG |
| 1fw13 | AAGGAGATATACATAATGGATAAGCCACCGTATTTACCAGTTTTCCGCCCCCCACGTCGCATTTACAACC |
| 1fw14 | AAGGAGATATACATAATGGATAAGCCACCGTATTTACCACGTCCCGTGCCCGTGCGTCGCATTTACAACC |
| 1fw15 | AAGGAGATATACATAATGGATAAGCCACCGTATTTACCACGTCCCCGCCCCCCATTATGGATTTACAACCGCTG |
| 1fw16 | AAGGAGATATACATAATGGATAAGCCACCGTATTTACCACGTCCCCGCCCCCCACGTCGCATTTACAACC |
| 1fw17 | AAGGAGATATACATAATGGATAAGCCACCGTATTTACCACGGCCCCGGCCCCCACGTCGCATTTACAACC |
| 1fw18 | AAGGAGATATACATAATGGATAAGCCACCGTATTTACCACGTCCCCGCCCTCCAAGACGCATTTACAACC |
| 1fw19 | AAGGAGATATACATAATGGATAAGCCACCGTATTTACCACGTCCCCGCCCCCCACGTCGCATTGGGAACA |
| 1fw20 | AAGGAGATATACATAATGGATAAGCCACCGTATTTACCACGTCCCCGCCGGGTGCGTCGCATTTACAACC |
| 1fw21 | AAGGAGATATACATAATGGATAAGCCACCGTATTTACCACGTACAATACCCCCACGTCGCATTTACAACC |
| 1fw22 | AAGGAGATATACATAATGGATAAGCCACCGTATTTACCACGTCCCCGCAATAGGCGTCGCATTTACAACC |
| 1fw23 | AAGGAGATATACATAATGGATAAGCCACCGTATGGTCCAGCTCCCCGCCCCCCATTGTGGATTTACAACCGCTG |
| 1fw24 | AAGGAGATATACATAATGGATAAGCCACCGTATTTACCACGTCCCCGCAGACGTCGTCGCATTTACAACC |
| 1rv1 | GGTGATGGTGGGATCTTATCAGCGGTTGTAAATGCGACGT |
| 1rv2 | GGTGATGGTGGGATCTTATCAGCGGTTGTAAATGCGCCCT |
| 1rv3 | GGTGATGGTGGGATCTTATCAGCGGGTGTAAATGCGACGT |
| 1rv4 | GGTGATGGTGGGATCTTATCAGCGGTTGTAAATGCGACGA |
| 1rv5 | GGTGATGGTGGGATCTTATCAGCGGTTACGAGGGCGACGT |
| 1rv6 | GGTGATGGTGGGATCTTATCAGCGGTTGTAAATCCTCCGT |
| 1rv7 | GGTGATGGTGGGATCTTATCAGCGGTTGTAAATCCATACT |
| 1rv8 | GGTGATGGTGGGATCTTATCAGCGGTTGTAAATGCGACGC |
| 1rv9 | GGTGATGGTGGGATCTTATCAGCGGTTGTAAATCCATAAT |
| 1rv10 | GGTGATGGTGGGATCTTATCAGCGGTTGTAAATGCGTCTT |
| 1rv11 | GGTGATGGTGGGATCTTATCATCTGTTCCCAATGCGACGT |
| 1rv12 | GGTGATGGTGGGATCTTATCAGCGGTTGTAAATCCACAAT |
| 2fw1 | AAGGAGATATACATATGGACAAACCACCGTACCTTCCGCGTCCTCGTCCGCCAAGACGTA |
| 2fw2 | AAGGAGATATACATATGGACAAACCCCCGTACCTTCCGCGTCCTCGTCCGCCAAGACGTA |
| 2fw3 | AAGGAGATATACATATGGACAAACCAACCCGCCTTCCGCGTCCTCGTCGGCCAAAACGTA |
| 2fw4 | AAGGAGATATACATATGGACAAACCAAACCGCCTTCCGCGTCCTCGTCCGCCAAGACGTA |
| 2fw5 | AAGGAGATATACATATGGACAAACCACCCTACCGTCCGCGTCCTCGTTACCGAAAACGTA |
| 2fw6 | AAGGAGATATACATATGGACAAAAGGCCCTACCGTCCGCGTCCTCGTCCGCCAAAACGTA |
| 2fw7 | AAGGAGATATACATATGGAAAAACCCACCCGCCTTCCGCGTCCTCGTCACCGAAAACGTA |
| 2fw8 | AAGGAGATATACATATGGACAAAAGGCCCTACCTTCCGCGTCCTCGTTTCCGAAAACGTA |
| 2fw9 | AAGGAGATATACATATGGACAAACCATGCCACCGTCCGCGTCCTCGTTACCGAAGACGTA |
| 2fw10 | AAGGAGATATACATATGGACAAACCACCGTGCCGTCCGCGTCCTCGTTGGCCAAGACGTA |
| 2fw11 | AAGGAGATATACATATGGACAAACCCTCCTACCTTCCGCGTCCTCGTCTCCCAAGACGTA |
| 2fw12 | AAGGAGATATACATATGGACAAACCCCCGTACCGTCCGCGTCCTCGTTGGCGAAGACGTA |
| 2fw13 | AAGGAGATATACATATGGACAAACACCCGCACCGTCCGCGTCCTCGTTACCGAAGACGTA |
| 2fw14 | AAGGAGATATACATATGGACAAAAGGCCGGTGCGTCCGCGTCCTCGTTTCCCAAGACGTA |
| 2rv1 | GTGATGGTGGGATCCTCATTAACGGTTGTAAATACGTCTTGGCGGACGAG |
| 2rv2 | GTGATGGTGGGATCCTCATTAACGGTTGTAAATACGTTTTGGCCGACGAG |
| 2rv3 | GTGATGGTGGGATCCTCATTAACGGTTGTAAATACGTTTTCGGTAACGAG |
| 2rv4 | GTGATGGTGGGATCCTCATTAACGGTTGTAAATACGTTTTGGCGGACGAG |
| 2rv5 | GTGATGGTGGGATCCTCATTAACGGTTGTAAATACGTTTTCGGTGACGAG |
| 2rv6 | GTGATGGTGGGATCCTCATTAACGGTTGTAAATACGTTTTCGGAAACGAG |
| 2rv7 | GTGATGGTGGGATCCTCATTAACGGTTGTAAATACGTCTTCGGTAACGAG |
| 2rv8 | GTGATGGTGGGATCCTCATTAACGGTTGTAAATACGTCTTGGCCAACGAG |
| 2rv9 | GTGATGGTGGGATCCTCATTAACGGTTGTAAATACGTCTTGGGAGACGAG |
| 2rv10 | GTGATGGTGGGATCCTCATTAACGGTTGTAAATACGTCTTCGCCAACGAG |
| 2rv11 | GTGATGGTGGGATCCTCATTAACGGTTGTAAATACGTCTTGGGAAACGAG |
